# Simulations reveal hybridization in Caribbean *Acropora* restoration poses low risk of genetic swamping but limited potential for adaptive introgression

**DOI:** 10.64898/2026.02.26.708281

**Authors:** Troy M. LaPolice, Colin N. Howe, Nicolas S. Locatelli, Christian D. Huber

## Abstract

Severe global declines in coral populations have driven growing demand for human intervention and restoration. One goal of restoration is to repopulate reef ecosystems through outplanting, which requires detailed understanding of target systems. However, long term ecological and reproductive data from interventions remain scarce. An exception to this are the critically endangered Caribbean corals, *Acropora palmata* and *A. cervicornis*, which have been central to restoration efforts in the region. These species serve as a unique case study due to the abundance of published data spanning ecology, and reproductive biology. In the wild, these species can cross to form an F_1_ hybrid, *A. prolifera*, though it is rarely used in restoration. It remains unclear whether *A. prolifera* is an evolutionary dead-end competing with its parents, or a potential bridge enabling genetic exchange via backcrossing. To evaluate benefits and risks of restoration among Caribbean *Acropora*, we developed a two-dimensional agent-based simulation using reproductive and ecological data to model realistic reef dynamics. Our model suggests the hybrid can facilitate introgression between parentals without outcompeting them. Yet, such introgression is too limited for large-scale or beneficial ancestry transfer except under ecologically unrealistic conditions or timescales significantly longer than those relevant for management. Thus, our model suggests that the risks of genetic swamping may be overstated, whereas hopes for adaptive introgression are also low, underscoring the value of simulations for generating long-term ecological and evolutionary insights relevant to coral restoration.

## Introduction

Worldwide, coral reefs continue to come under ever-increasing threats from disease and anthropogenic forces^1^. Each year, global ocean temperatures continue to rise, leading to unprecedented—and accelerating—coral death^2,3^. Climate-driven stressors pose a particularly severe threat to coral reefs, which are foundational to marine ecosystems. By offering both structural habitat and food resources, reefs sustain extraordinary biodiversity, with estimates indicating that at least 25% of marine species depend on these reefs^4,5^. Further, one study predicts that without corals, global tropical fish biodiversity could decline by approximately one half^6^. Such a decline would trigger cascading effects throughout marine ecosystems, and disrupt the many human and ecological communities that rely on these ecosystem services. Additionally, corals are not just important for biodiversity, but also provide coastline protection by dissipating potentially damaging wave energy before it can make landfall. Due to their importance in the ocean ecosystem—as well as the coastline protection corals provide^7,8^—reefs are estimated to provide a benefit to humans worth $352,000, per hectare of reef, per year^8^. As such, the demand for human intervention to restore decimated coral populations has grown immensely^1,9^.

As coral populations continue to decline, governments, local, and indigenous communities are under increasing pressure to implement the most effective and efficient restoration strategies to combat mass coral mortality. Ocean-based coral restoration programs propagate coral fragments that are grown *in situ* on nursery structures. These fragments may also be sourced from the wild, including “fragments of opportunity” generated by boat groundings^10^, naturally detached pieces, or fragments intentionally collected from live colonies^11^. These fragments are then planted directly on the benthic substrate—with the goal of maturation alongside other planted fragments^12–14^. Because this method (called outplanting) focuses on degraded reefscapes, requires human selection of colonies, and manual planting of fragments on the reef, restoration practitioners have potentially significant influence in determining the resulting restored reef architecture. Thus, it is important to understand the long term impact of different outplanting schemes on population genetics and species dynamics within a restored reef.

Coral restoration programs established in the Atlantic-Caribbean region have found success incorporating two critically endangered coral species, *Acropora cervicornis* and *A. palmata,* into restoration programs for over 20 years^13–17^. These two Caribbean *Acropora* species can also naturally hybridize to produce a developmentally viable F_1_ hybrid, *A. prolifera*^18,19^. Further, there is evidence that *A. prolifera* may rarely backcross with its parental species^19^.

Although *A. prolifera* is rare at the ecosystem level—for example, among 1,676 *Acropora* observations in the 20-year Florida Fish and Wildlife Conservation Commission Disturbance Response Monitoring program, only 9 were *A. prolifera* (compared to 1,597 *A. cervicornis* and 70 *A. palmata*^20^)—several populations of *A. prolifera* exhibit high genotypic diversity^21,22^. In some Caribbean locations, these hybrid populations have reached relative abundances comparable to those of the parental taxa^23,24^.These findings are consistent with earlier studies showing that the hybrid maintains long-term survival and has shown similar ecological viability (i.e., *in situ* mortality, disease and predation tolerances, and susceptibility to algae overgrowth) to its parental species^25,26^. Although subsequent research has identified variation in growth rates and responses to environmental stressors (e.g., predation) among distinct hybrid genotypes, no evidence suggests reduced hybrid survival across different coral restoration approaches^27,28^. Thus, despite mounting evidence that *A. prolifera* could be a valuable addition to coral restoration programs, the introduction of hybrid taxa to conservation efforts has remained limited and its usage may require scrutiny to maintain species and genetic diversity of the parental species.

The long term consequences of active restoration and the impacts on gene flow continue to be the source of great debate, leading researchers to propose different theories for the result of introgression^29–33^. Due to the propensity for hybridization within the *Acropora* genus^32,34^, some worry that introgression may be damaging to the population, with concern that widespread introgression could lead to “genetic swamping^32^”. This process may blur genomic boundaries between species, potentially eroding local adaptation and disrupting existing population dynamics. It can also lead to the loss of valuable genetic diversity^32,35^, or reduce species diversity in cases of “demographic swamping.” An additional concern regarding hybridization in *Acropora* is the potential for outbreeding depression—potentially resulting from the hybrid facilitating maladaptive or deleterious combinations of genetic variants^30,31,34^. Concerns of outbreeding depression or genetic swamping may be alleviated if the F_2_s and/or backcrosses exhibit hybrid breakdown^36^, limiting the ability of *A. prolifera* to serve as a vehicle for introgression. Even without backcrossing or hybrid breakdown, a reduction in species diversity could also occur if hybrid vigor allows a functionally sterile hybrid to outcompete parentals^29,37,38^.

Contrary to these concerns, others have argued hybridization could, instead, facilitate higher rates of heterozygosity, increase standing genetic variation, and introduce beneficial alleles through adaptive introgression. This could result in increased genetic diversity which could improve adaptability, and be beneficial for restoration^33,38,39^. Concerns about the uncertain impacts of hybridization have led practitioners to debate and often limit the use of *A. prolifera* in restoration contexts.

Understanding these impacts requires long-term field monitoring paired with genomic analysis. However, restoration programs are ultimately limited by monetary cost, staffing, and time to collect this data^40^. As a consequence, the data on a majority of post outplanting monitoring projects do not extend beyond 18 months^1^. Further, identifying how human selective restoration influences the community structure, and rates of introgression at restored sites, remains a challenge due to long generation times of corals^41^.

Given the urgency to act in response to rising ocean temperatures, a biologically realistic simulation provides an efficient framework to evaluate both adaptive and evolutionary processes. Simulation studies provide the ability to test multiple outplanting regimes, without spending prohibitive amounts of money, or removing fragments from the limited remaining healthy coral populations. Here, we utilize an agent-based, two-dimensional simulation model which allows for rapid genetic monitoring of various outplanting regimes.

Agent-based simulations model complex behaviors by simulating many individual behaviors of independent units or “agents” within the simulation. First applied in biology to model social structure in bee colonies^42^, this modeling of the microscale allows for an accurate reconstruction of the larger picture. Early work on corals by Sleeman et al. 2005^43^ used an agent-based spatial simulation to explore how different outplanting arrangements influence coral cover and structural complexity, showing that evenly spaced transplants of fast-growing corals can accelerate reef recovery. More recently, Carturan et al. 2020^44^ developed a trait-based agent model to evaluate how variation in coral functional traits and local interactions shapes community trajectories, offering a framework for exploring ecological dynamics using empirically informed simulations. These ecological models advanced restoration-relevant questions but did not incorporate genetic or evolutionary processes. Evolutionary modeling efforts by Matz et al., (2018, 2020)^45,46^ have used agent-based simulations to evaluate how standing genetic variation and population connectivity influence coral responses to warming, highlighting the importance of evolutionary processes in shaping long-term resilience. The 2020 Matz et al., study also noted that understanding patterns of gene flow can help inform the principles behind assisted gene flow and related interventions, but did not evaluate specific restoration actions^46^.

Here, we apply an interdisciplinary agent-based approach to the critically endangered *Acropora* system. By integrating empirical life history traits and genomic data with agent-based simulations, we model individual coral genomes in SLiM^47^ to investigate gene flow, assess the potential for adaptive introgression, and identify the conditions under which swamping may occur. We find that following a restoration attempt, a major factor in dictating long term species dynamics is the initial outplanting species ratio. Further, we find that there is minimal concern for genetic swamping in parental species due to low introgression rates in the Caribbean *Acropora* system. Consequently, the potential for adaptive introgression is also low.

## Results

We used computational modeling to better understand three key aspects of the Caribbean *Acropora* system that are directly relevant when applying active restoration efforts. First, we evaluated whether varying levels of gametic incompatibility between Caribbean *Acropora* species influence gene flow, and test how different assumptions regarding the directionality of gene flow, backcrossing, and F_2_ production affect the extent of introgression in each species. Second, we examined the impact of various parental outplanting schemes on introgression rates and species dynamics— including the potential for genetic or demographic swamping of parental species. Finally, we assessed whether backcrossing can facilitate adaptive introgression by enabling the transfer of advantageous alleles between species.

### Agent-based simulation framework

We developed an agent-based model in SLiM (v4.0)^47^ to examine how restoration strategies and assumptions about reproductive isolation affect introgression and the population dynamics of *A. cervicornis*, *A. palmata,* and *A. prolifera* in simulated restored sites over ecological and evolutionary timescales. The model represents a two-dimensional reef environment in which individual corals occupy spatial locations and interact locally. The SLiM^47^ simulation allows us to model overlapping generations, reflecting that corals are long-lived organisms in which adults, juveniles, and newly settled recruits coexist and experience different life-history processes simultaneously. Lastly, all individuals in our model have simulated diploid genomes, allowing population ancestry to be tracked through time (Fig. 1). Together, these features enable us to examine demographic and evolutionary dynamics across multiple temporal horizons. To facilitate interpretation, we therefore structure the results around three predefined timescales—200 years (human-relevant), 1,000 years (long-term ecological), and, in a reduced subset of simulations, 6,000–20,000 years (evolutionary)—which provide complementary perspectives on short-, intermediate-, and deep-time processes.

**Figure 1:**
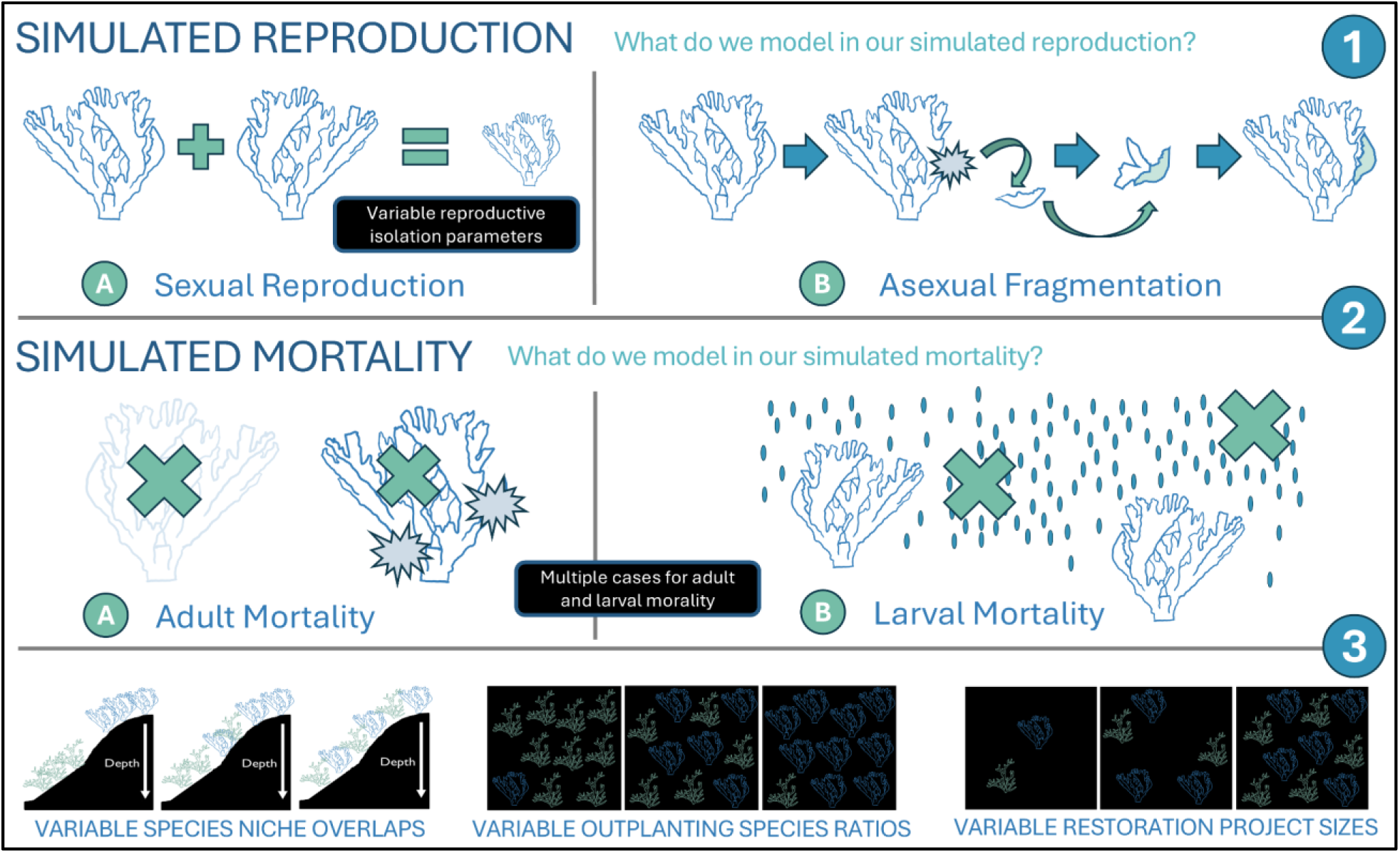
Overview of our computational agent based model. Our model simulates multiple realistic modes of reproduction and mortality. (1) We replicate sexual reproduction (under a variety of different reproductive isolation scenarios which we explore in this present paper), as well as asexual clonal fragmentation-based reproduction. (2) We simulate multiple types of mortality imposed at both the larval and adult stage (see Methods). (3) Finally, we model variable outplanting schemes and intrinsic ecological dynamics to assess their impact on species dynamics and gene flow.

To accurately model reproduction, we incorporated empirical data from Fogarty et al. (2012) and explored a broad parameter space for aspects of genetic isolation that remain uncertain or poorly characterized. Another important aspect of modeling coral biology is the ability to reproduce asexually. Asexual reproduction occurs through clonal propagation, producing genetically identical fragments that establish nearby. Together, these processes allow the model to reflect the mixed reproductive strategy characteristic of branching corals (Fig. 1).

Mortality occurs across multiple life-history stages in our model, with adult colonies experiencing background mortality that, in some simulations, was elevated episodically to simulate disturbance events such as bleaching (Fig. 1). We assume larvae experience high stochastic mortality (Fig. 1), with further settlement success rates based on the local population densities inferred from empirical studies of genotypic and clonal diversity in *A. cervicornis*^48^.

We simulate under a variety of conditions to determine how the system responds under different restoration practices. To explore the role of ecological context, we examined outcomes under varying degrees of spatial competition among *Acropora* species. Empirical evidence indicates that *A. palmata* and *A. cervicornis* occupy partially differentiated niches along a depth gradient, with *A. palmata* more common at shallow depths but otherwise broadly sympatric^49–51^. We evaluated outcomes across a range of species niche competition scenarios to determine the influence on introgression and system behavior.

We refer to these competition scenarios as “niche overlap.” The niche overlap can be thought of as representing a depth gradient from monotypic outplanting to mixed-species schemes where the degree of interspecies competition along the gradient is determined by a niche overlap parameter. Similarly, we explored different outplanting species ratios where *A. cervicornis* and *A. palmata* are planted in both even and biased ratios to assess how outplanting influences long-term introgression and species dynamics of the system. Lastly, we also evaluated how project size (measured as the total number of outplanted individuals) shapes long-term outcomes and variability (Fig. 1).

### Reproductive isolation has limited effects on species dynamics

To address our first aim, we examined how *A. prolifera* gamete viability influences species ratios following outplanting. We initialized simulations with equal numbers of each parental taxon and ran them for 1,000 years under baseline parameters for reproduction, settlement, and mortality. This baseline assumes *A. palmata* or *A. cervicornis* eggs accept heterospecific sperm at a rate based on mixed-sperm laboratory trials reported by Fogarty et al.^52^, and that 1% of larvae survive initially and have a chance to settle (depending on local density). We next introduce a bias parameter (γ) to control the viability of hybrid gametes (a prezygotic form of hybrid breakdown) relative to the parentals (see Methods), reflecting the possibility that gametes may be less viable due to asymmetrical recombination in hybrids^53–55^. When γ = 0, hybrid gametes are inviable, and cannot produce offspring; when γ = 1.0, they are as viable as those of the parentals. We tested five γ values (0, 0.25, 0.5, 0.75, 1.0) to assess how varying levels of gametic viability influence species ratios through time (Fig. 2A–B).

**Figure 2:**
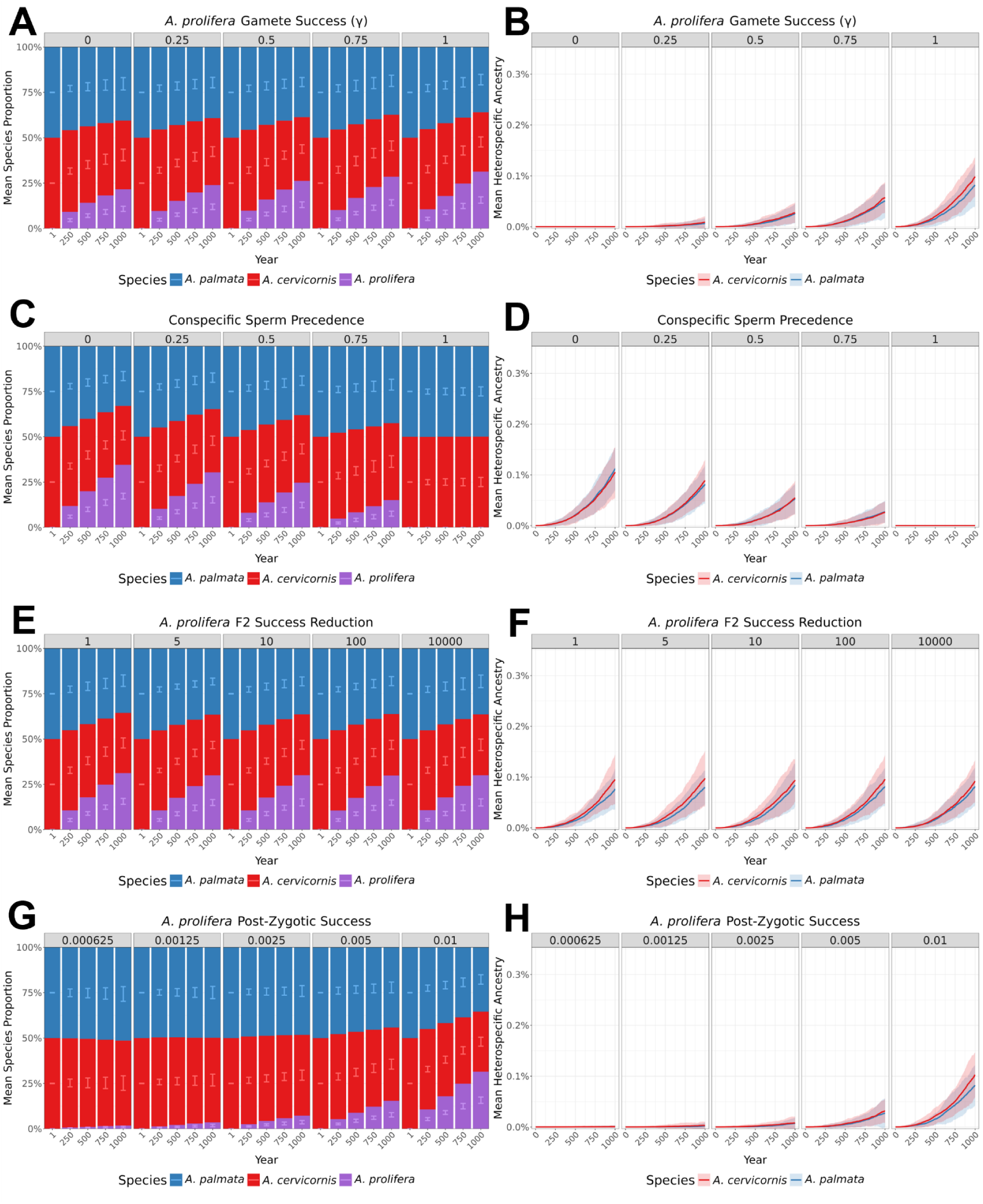
Effects of reproductive parameters on species composition and ancestry through time. Panels (A, C, E, G) show species ratios, and panels (B, D, F, H) show mean heterospecific ancestry, each faceted by a different parameter: hybrid gamete success (γ), conspecific sperm precedence (CSP), F_2_ success reduction where F_2_ viability was reduced 5×, 10×, 100×, or 1,000× relative to the baseline initial larval survival of 1%, and *A. prolifera* larval success where we halved all hybrid larval survival in four steps (0.5%, 0.25%, 0.125%, 0.0625%). For all species-ratio panels (A, C, E, G), the x-axis represents snapshots of simulation years and the y-axis shows the mean species ratio across replicates. Stacked bars display the mean proportion of each taxon, with error bars indicating ±2× standard deviation between replicates (n=50). Colors: Blue = *A. palmata*, red = *A. cervicornis*, purple = hybrid *A. prolifera*. For all heterospecific-ancestry panels (B, D, F, H), solid lines depict mean heterospecific ancestry across replicates (n=50) and shaded areas show ±2× the standard deviation between replicates. Colors: blue = *A. palmata*, red = *A. cervicornis*.

The effect of γ on introgression and species dynamics was limited, particularly over short restoration timescales (first 200 years, Supplementary Figs. 1A–B). Species trajectories remained largely consistent across γ values (Fig. 2A; Supplementary Fig. 1A), with the primary impact seen in the number of backcrossed *A. prolifera* produced (as backcrossing requires viable *A. prolifera* gametes) (Supplementary Fig. 2). A similar trend was observed for F_2_ individuals (Supplementary Fig. 2). Although population-level species ratios changed little, patterns of introgression were more directly shaped by hybrid gamete viability. When hybrid gametes were inviable (γ = 0.0), introgression ceased entirely (Fig. 2B). As γ increases, backcrossing and introgression rose slightly but remained minimal genome-wide (Fig. 2B), particularly over short restoration timescales where the trend was negligible (Supplementary Fig. 1B).

Because F_1_ hybrids form directly from parental gametes, γ does not influence their production. To examine another form of reproductive isolation that could limit hybridization, we introduced a parameter for conspecific sperm precedence (CSP). CSP determines the likelihood that *A. palmata* or *A. cervicornis* eggs accept heterospecific sperm, ranging from no preference (CSP = 0) to complete conspecific exclusivity (CSP = 1). Baseline CSP values were set to 0.333 for *A. palmata*, and 0.03 for *A. cervicornis*, based on mixed-sperm laboratory trials from Fogarty et al., (2012)^52^. However, to explore a broader range of potential gametic incompatibilities, we also tested a range of symmetrical CSP values (Figs. 2C–D; Supplementary Figs. 1C–D; Supplementary Fig. 3) for both species (0, 0.25, 0.5, 0.75, 1.0).

Compared with the γ parameter, F_1_ formation is more strongly influenced by conspecific sperm precedence (CSP), as hybrid production requires an egg from one parental species to be fertilized by heterospecific sperm. As a result, increasing CSP reduced the number of F_1_ hybrids and exerted a somewhat greater effect on overall species dynamics (Fig. 2C, Supplementary Fig. 3). Although CSP also influences the degree of backcrossing, its effect on the backcrossing rate was weaker than that of γ (Supplementary Fig. 2; Supplementary Fig. 3). This difference arises because γ reduces the viability of both *A. prolifera* gamete types, whereas CSP operates only through fertilization bias on the egg side. Nonetheless, the overall pattern of introgression remains comparable to that observed under varying γ values, showing minimal impact, particularly over shorter time scales (Fig. 2D; Supplementary Fig. 1D).

CSP exerts the strongest impact on hybridization when both species have complete conspecific sperm precedence (CSP = 1). Under this scenario, eggs can be fertilized only by conspecific sperm, preventing F_1_ formation and backcrossing altogether (Fig. 2C; Supplementary Fig. 1C; Supplementary Fig. 3). However, as soon as one species’ eggs retain partial compatibility with heterospecific sperm (CSP < 1), hybridization and gene flow resumes, allowing bidirectional exchange even under asymmetric isolation. I.e., this allows for bidirectional gene flow between the species, even when one species’ eggs can only form zygotes with conspecific sperm. We next tested another reproductive isolation mechanism: reduced postzygotic hybrid viability in *A. prolifera* (Figs. 2E–H).

### Hybrid viability strongly influences hybrid persistence

We examined postzygotic isolation via reduced hybrid viability using two simulation designs. In the first, F_1_s formed normally but produced F_2_s with reduced success. F_2_ viability was reduced 5×, 10×, 100×, or 1,000× relative to the baseline initial larval survival of 1% (Figs. 2E–F; Supplementary Figs. 4A–B; Supplementary Fig. 5). In the second, all hybrid larvae (F_1_s, F_2_s, and backcrosses) experienced reduced fitness (Figs. 2G–H; Supplementary Figs. 4C–D; Supplementary Fig. 6). Using our baseline larval survival of 1%, we halved hybrid survival in four steps (0.5%, 0.25%, 0.125%, 0.0625%) to evaluate population outcomes (Figs. 2G–H; Supplementary Figs. 4C–D; Supplementary Fig. 6).

In the first reduced-hybrid-success model, where F_2_s exhibit reduced success, the effect on overall species ratios and introgression rates was negligible (Fig. 2E–F). This pattern held across both short- and long-term timescales (Supplementary Figs. 4A–B). The only measurable effect was a reduction in the number of F_2_ individuals, with zero F_2_ individuals produced under higher levels of F_2_ success reduction (Supplementary Fig. 5). In contrast, when reduced viability was applied to all hybrid types, the impact on population dynamics was substantial. Overall introgression declined sharply, and species ratios shifted toward lower *A. prolifera* abundance across both short- and long-term timescales (Supplementary Figs. 4C–D; Figs. 2G–H; Supplementary Fig 6).

Together, these results demonstrate that F_2_ success reduction alone exerts little influence on species ratios, whereas even moderate reductions in hybrid (F1s, F2s, and backcrosses) larval viability markedly constrain introgression and the persistence of *A. prolifera*. Thus, postzygotic isolation through reduced hybrid larval fitness can substantially limit gene flow between parental species. Having established that hybrid fitness strongly shapes long-term species dynamics, we next examined how restoration strategies influence these outcomes.

### Restoration practices alter species balance and introgression

To examine how restoration design influences long-term species outcomes, we simulated several restoration scenarios that varied in outplanting ratios (Fig. 3A–B; Supplementary Figs. 7A–B; Supplementary Fig. 8), niche overlap (Fig. 3C–D; Supplementary Figs. 7C–D; Supplementary Fig. 9) and project size. We first modeled biased outplanting, where one parental species was overrepresented at the start of the restoration. We simulated five initial *A. palmata* to *A. cervicornis* outplanting ratios (0.1, 0.25, 0.5, 0.75, and 0.9), and assessed their effects on species dynamics and introgression. We found that species dynamics are strongly impacted by a skewed outplanting ratio (Fig. 3A; Supplementary Fig. 7A; Supplementary Fig. 8). Prioritizing a single species when outplanting can drive long term dominance of that species within the reef (Fig. 3A; Supplementary Fig. 8B).

**Figure 3:**
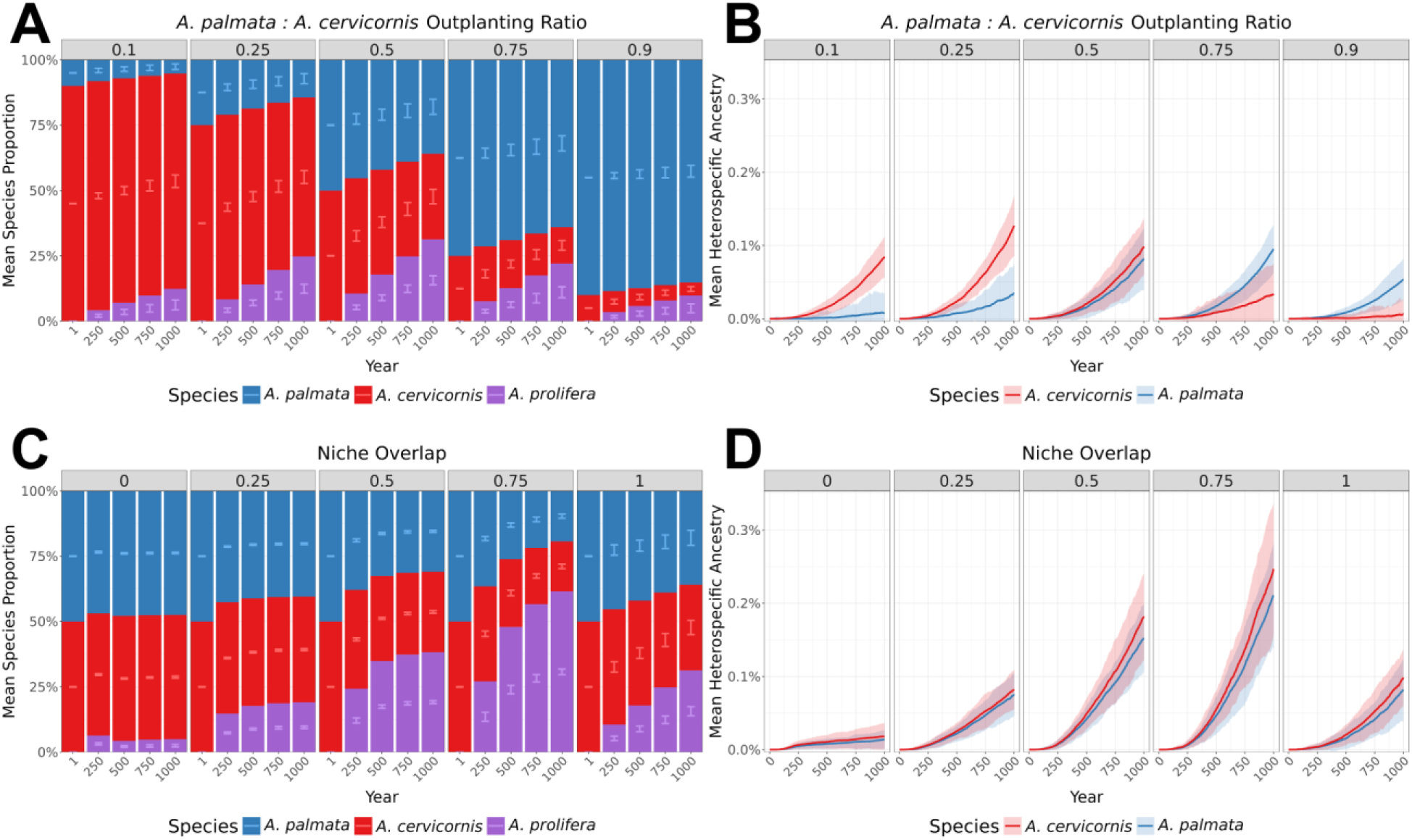
Effect of initial outplanting ratio and niche overlap on introgression and species dynamics. Species ratios (A, C) and mean heterospecific ancestry (B, D) through time under two conditions: varying initial outplanting ratios of *A. palmata* to *A. cervicornis* where a ratio of 0.1 means 10%:90% *A. palmata:A. cervicornis* and 0.9 means 90%:10% *A. palmata:A. cervicornis* (A, B), and varying degrees of niche overlap where 0 = no overlap and 1 = full overlap (C, D). For species-ratio panels (A, C), the x-axis represents snapshots of simulation years and the y-axis shows the mean species ratio across replicates. Stacked bars display the mean proportion of each taxon, with error bars indicating ±2× standard deviation between replicates (n=50). Colors: blue = *A. palmata*, red = *A. cervicornis*, purple = hybrid *A. prolifera*. For heterospecific-ancestry panels (B, D), solid lines depict mean heterospecific ancestry across replicates (n=50) and shaded areas show ±2× the standard deviation between replicates. Colors: blue = *A. palmata*, red = *A. cervicornis*.

This skew also influenced introgression patterns, which remained negligible at first (Supplementary Fig. 7B), but increased after approximately 200 years (Fig. 3B). Most backcrossed individuals carried ancestry proportions of roughly 75% from the majority taxon and 25% from the minority (Supplementary Fig. 8), representing first-generation backcrosses (BC_1_s). Consequently, introgression occurred primarily from the minority toward the majority species, reflecting asymmetric backcrossing opportunities driven by imbalance in relative abundance (Fig. 3B).

In summary, when outplanting was heavily skewed toward one taxon, that species maintained dominance on the reef over hundreds of years, often suppressing or excluding the minority parental species. Such outcomes indicate that unequal outplanting can produce lasting imbalances in community composition unless the minority taxon occupies an independent ecological niche. To test this possibility, we next examined how niche overlap modulates competition and hybridization dynamics.

With fully independent niches (e.g., depth-separated habitats), each parental species reached its carrying capacity within its own niche, leaving little opportunity for hybrid establishment and thus a low abundance of *A. prolifera* (Fig. 3C). Conversely, under partial niche overlap, all three taxa persisted for centuries (Fig. 3C). Interestingly, *A. prolifera* is most successful with high, but not complete niche overlap (i.e., 50-75%), eventually dominating the species composition (Fig. 3C; Supplementary Fig. 9). Introgression remained minimal on restoration timescales (< 200 years) but increased modestly over longer periods, peaking under partial but high niche overlap (Fig. 3D; Supplementary Fig. 7D).

We then assessed how niche structure interacts with biased outplanting. When niches partially overlapped, the favored outplanted species expanded rapidly at first, taking over both its preferred niche and the intermediate habitat where the species’ ranges met (Fig. 4). However, as the other parental species gradually grew into its own niche, it began competing with the dominant taxon in this shared zone, leading to the formation of more hybrids within this hybrid zone. These new hybrids were then able to colonize both parental and the intermediate habitats. Similar to other scenarios, these dynamics unfolded slowly, with noticeable effects emerging only after about 200 years (Fig. 4; Supplementary Fig. 10).

**Figure 4:**
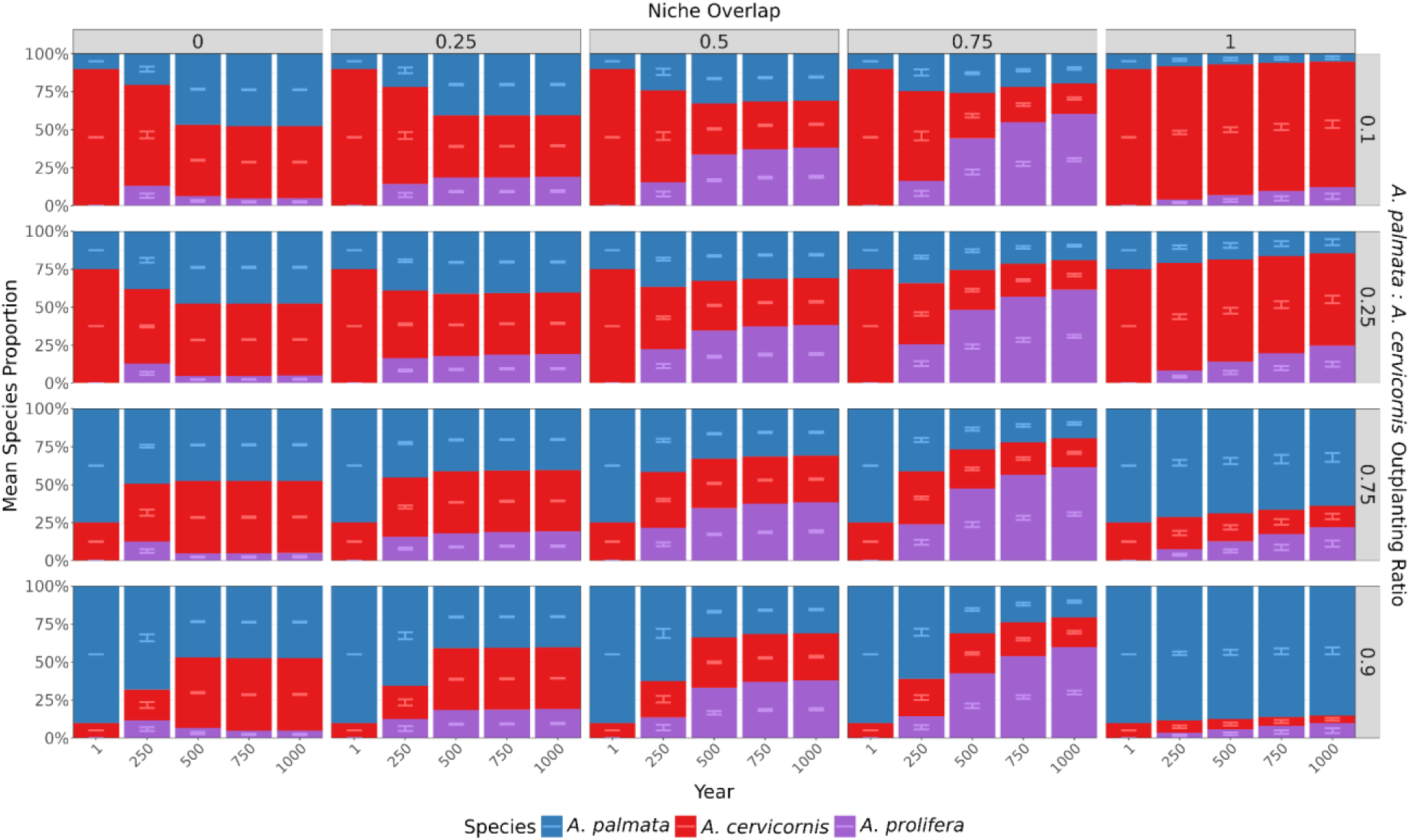
Effect of the interaction between initial outplanting ratio and niche overlap on species dynamics. Species ratios through time, faceted the degree of niche overlap where 0 = no overlap and 1 = full overlap (top) and the initial outplanting ratio of *A. palmata* to *A. cervicornis* (right), and the initial outplanting ratio of *A. palmata* to *A. cervicornis* (right) where a ratio of 0.1 means 10%:90% *A. palmata:A. cervicornis* and 0.9 means 90%:10% *A. palmata:A. Cervicornis*. The x-axis represents snapshots of simulation years and the y-axis shows the mean species ratio across replicates. Stacked bars display the mean proportion of each taxon, with error bars indicating ±2× standard deviation between replicates (n=50). Colors: blue = *A. palmata*, red = *A. cervicornis*, purple = hybrid *A. prolifera*.

Finally, the initial outplanting size is an important consideration in restoration design, as practitioners must allocate limited resources effectively. To evaluate how outplanting size influences long-term outcomes, we tested varying starting sizes and their effects on species dynamics and introgression (Fig. 5; Supplementary Fig. 11). We found that average species ratios and introgression levels remained similar on average across different initial outplant sizes, but the variability among replicates changed markedly. Larger outplanting size produced more consistent outcomes across 150 replicate simulations than smaller size (see errorbars Fig. 5A, Supplementary Fig. 11A; shading Fig. 5B, Supplementary Fig. 11B). Assuming a closed system with a spatial extent similar to our simulated space, outplanting at least a few hundred individuals is required for consistent and predictable outcomes. This effect was most apparent after approximately 200-250 years where we see replicates begin to diverge from one another (see Fig. 5B; Supplementary Fig. 11B). Interestingly, we observed slightly fewer hybrids on average when the initial outplant size was small (Supplementary Fig. 12), particularly early on (Supplementary Fig. 12A).

**Figure 5:**
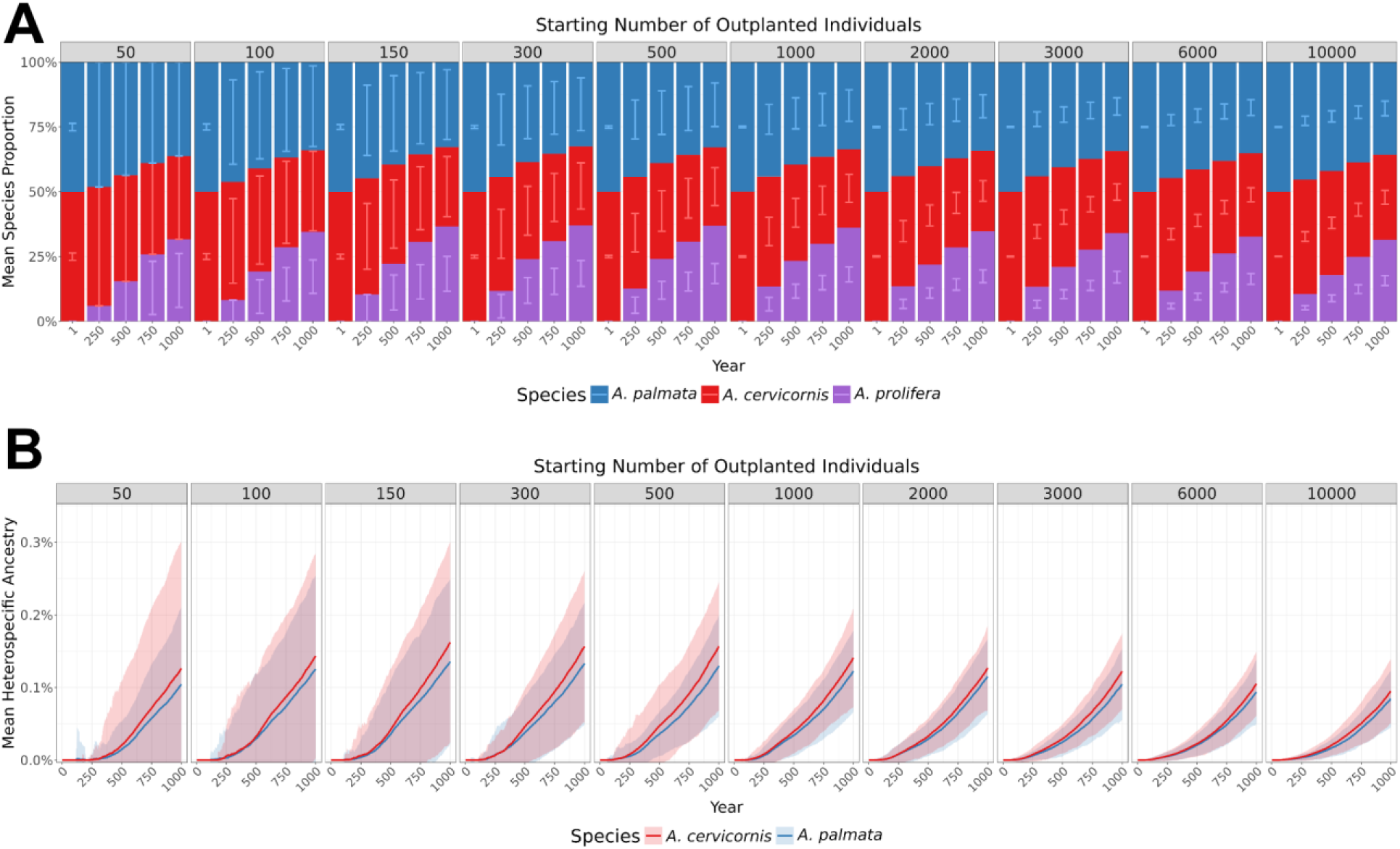
Effect of outplant project size on restoration outcomes. (A) Species ratios through time, faceted the number of initially outplanted individuals, the x-axis represents snapshots of simulation years and the y-axis shows the mean species ratio across replicates. Stacked bars display the mean proportion of each taxon, with error bars indicating ±2× standard deviation between replicates (n=50). Colors: blue = *A. palmata*, red = *A. cervicornis*, purple = hybrid *A. prolifera*. (B) solid lines depict mean heterospecific ancestry across replicates (n=50) and shaded areas show ±2× the standard deviation between replicates. Colors: blue = *A. palmata*, red = *A. cervicornis*.

Together, these findings indicate that both restoration design—especially outplanting ratios and project scale—and intrinsic ecological factors such as niche overlap strongly shape long-term species balance and hybridization dynamics. Having characterized the ecological, demographic and genetic context of introgression, we next examined whether hybridization could facilitate adaptive introgression under environmental selection.

### Adaptive introgression is limited by timescale

Hybrid taxa have been proposed as potential conduits for transferring adaptive alleles, such as variants conferring heat tolerance, between parental species^32–34^. To test this, we introduced a single beneficial allele into all outplanted *A. palmata* individuals. In our simulations, this hypothetical allele protects against simulated bleaching events which increase in frequency over time, consistent with projected future climate scenarios for coral reefs^56,57^. We modeled the spread of this resistance allele under varying selective strengths, where carrying the allele reduced the probability of adult mortality during bleaching events by 1%, 5%, 10%, 25%, 50%, 75%, or 90%. Individuals homozygous for the allele experienced the full benefit, while heterozygotes received half that advantage, corresponding to an additive fitness model.

We found that the donor species (i.e., the parental species initially carrying the beneficial allele), along with hybrids that inherited one copy, rapidly outcompeted the other parental species before enough time passed for introgression to occur (Fig. 6). Consequently, the potential for adaptive introgression in this system is limited by the timescale required for gene flow to take effect.

**Figure 6:**
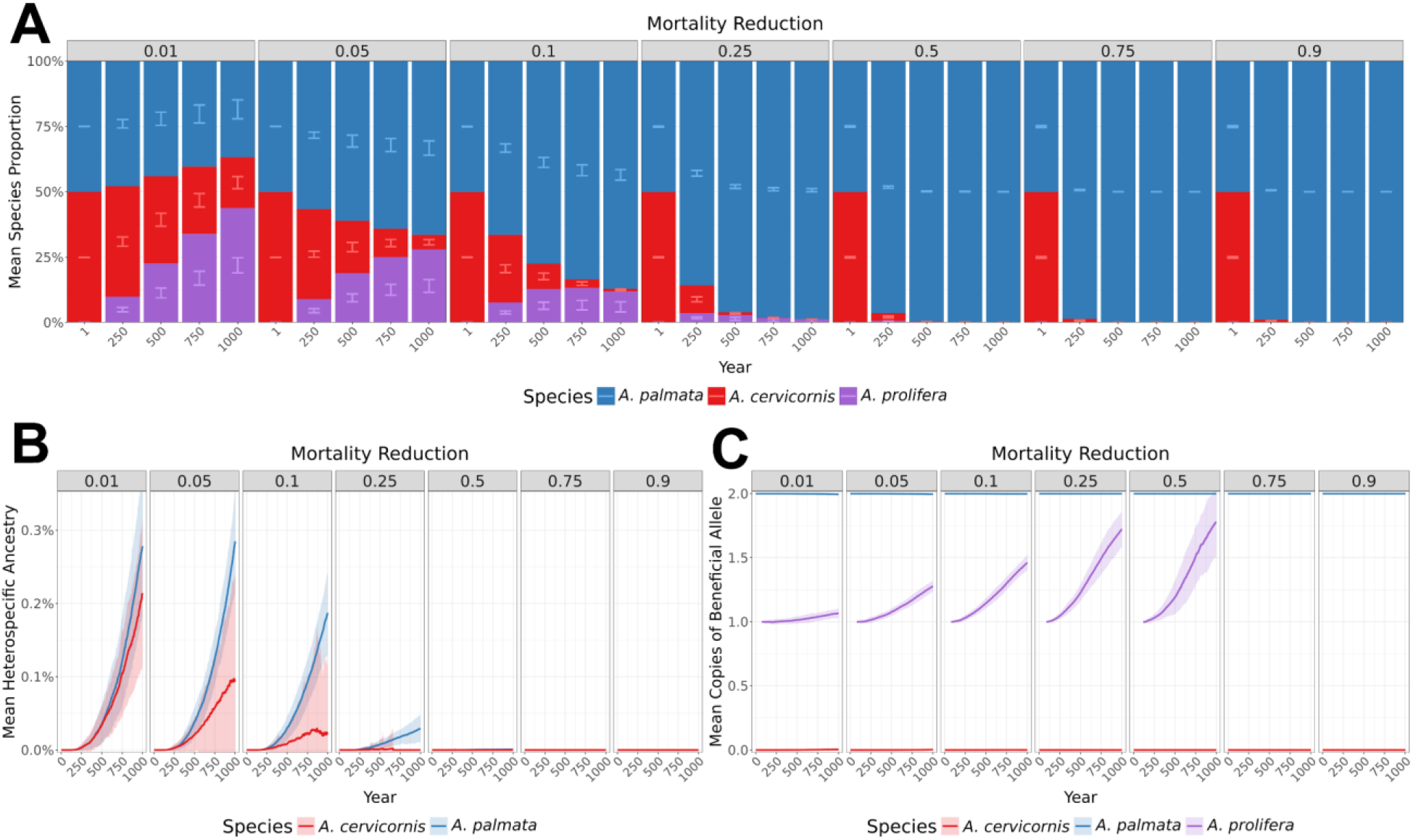
Effect of single additive beneficial locus initially fixed in *A. palmata* on introgression and species dynamics when niches fully overlap. (A) Species ratios through time, faceted by the mortality reduction from carrying an additive beneficial allele in *A. palmata*. The x-axis represents snapshots of simulation years and the y-axis shows the mean species ratio across replicates. Stacked bars display the mean proportion of each taxon, with error bars indicating ±2× standard deviation between replicates (n=50). Colors: blue = *A. palmata*, red = *A. cervicornis*, purple = hybrid *A. prolifera*. (B) Mean heterospecific ancestry through time, faceted the mortality reduction from carrying the beneficial allele. The solid lines depict the mean heterospecific ancestry in the population across replicates and shaded areas show ±2× the standard deviation between replicates (n=50). Blue lines and shading represent *A. palmata*, red lines and shading represent *A. cervicornis.* (C) The mean number of copies of the beneficial allele in the three Acroporids through time, faceted by the mortality reduction. Blue lines and shading represent *A. palmata*, red lines and shading represent *A. cervicornis*, purple represents *A. prolifera.* Shaded areas show ±2× the standard deviation between replicates (n=50).

Interestingly, introgression rates between parentals were highest when the selective advantage of the beneficial allele was weak (Fig. 6B). Importantly, there was a modest impact on the species dynamics under shorter timescales (Supplementary Figs 13A; 14A) and this introgression was not seen until more than 200 years into the simulation (Fig. 6; Supplementary Figs. 13B–C). Slower competitive exclusion under weak selection extended coexistence between species, increasing opportunities for gene flow and backcrossing (Fig. 6B, Supplementary Fig 14). When the selective advantage was strong, then introgression ceased entirely as hybrids were not generated due to the rapid die off of the species without the selective loci (Fig. 6C; Supplementary Fig. 13C). When examining the direction of introgression, we found that gene flow was biased toward the parental species carrying the beneficial allele, *A. palmata* (blue line, Fig. 6B). This asymmetry likely reflects demographic imbalance, because *A. palmata* was more abundant (due to its higher fitness from carrying the beneficial allele), and hybrids more frequently backcrossed into this species, producing greater gene flow in that direction, similar to the pattern observed under skewed outplanting ratios. Introgression into the other parental species was limited, and the beneficial allele did not reach detectable frequencies in *A. cervicornis*. Instead, its frequency slowly increased within the hybrid population through repeated backcrossing with *A. palmata* which allowed the backcrossed hybrids a chance to obtain both beneficial *A. palmata* alleles over longer timescales (Fig. 6C; Supplementary Fig 13C).

We next tested how niche overlap affects adaptive introgression. Independent niches could let the non-adapted species (in this case *A. cervicornis*) to persist longer, extending the window for gene flow. Using the same additive model (1–90% mortality reduction), we simulated independent and 50% overlapping niches (Fig. 7).

**Figure 7:**
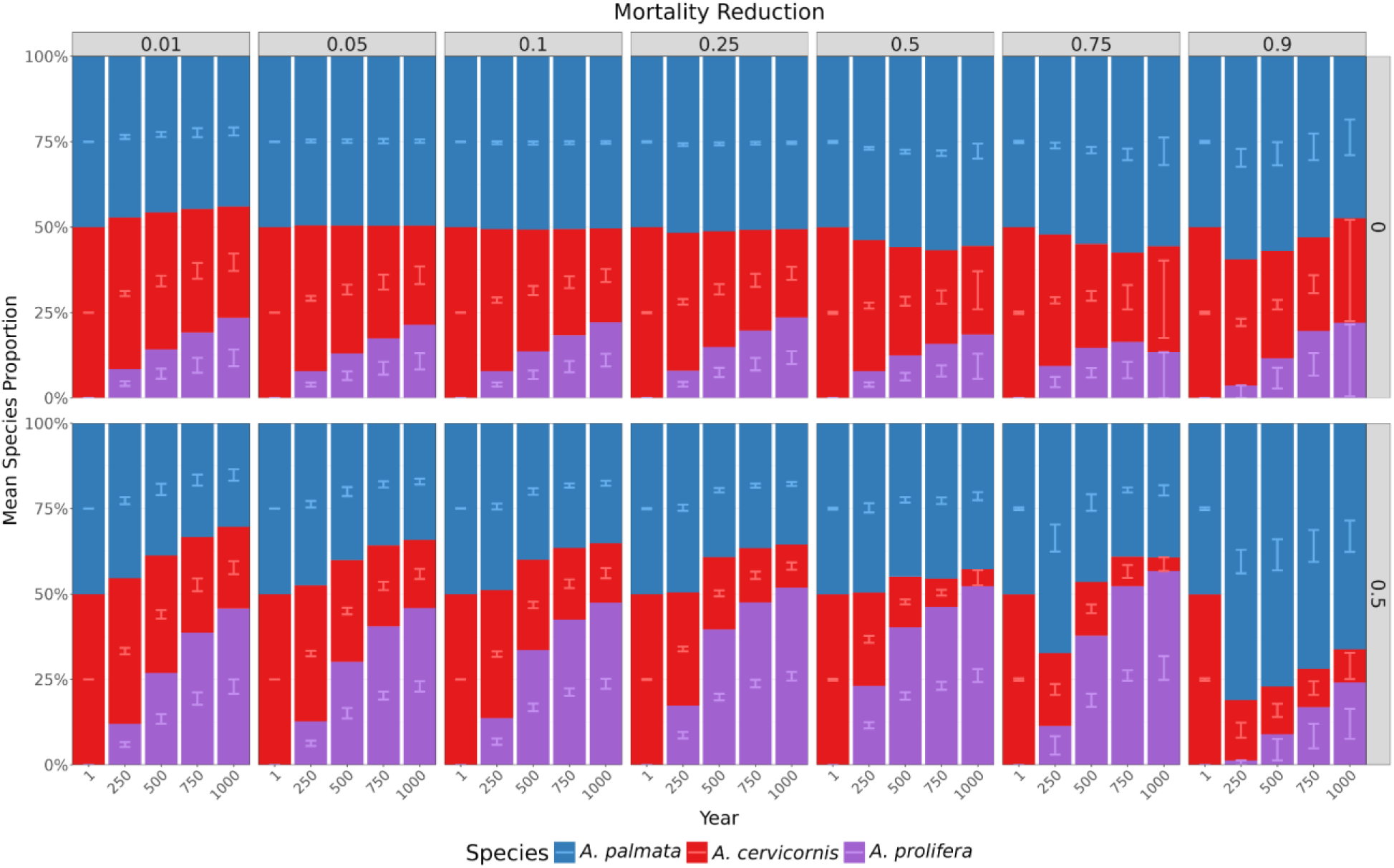
Effect of variable niche overlaps and single additive beneficial locus initially fixed in *A. palmata* on species dynamics. (A) Species ratios through time, faceted the mortality reduction from carrying the beneficial allele (top) and the niche overlap where 0 = no overlap and 1 = full overlap (right). X-axis represents snapshots of the mean species ratio (y-axis) in different years through the simulation. The stacked bars represent a mean of the species ratio in the population across replicates. The error bars represent ±2× the standard deviation between replicates (n=50). Colors: blue = *A. palmata*, red = *A. cervicornis*, purple = hybrid *A. prolifera*.

With partial or independent niches, *A. cervicornis* (which lacked the hypothetical beneficial allele) indeed persisted longer and reached larger populations than under complete overlap. Its greatest abundance occurred when niches were independent, allowing coexistence even under strong selection favoring *A. palmata* (Fig. 7). This persistence prolonged coexistence, slightly increasing opportunities for adaptive introgression (Fig. 8; Supplementary Fig. 15). However, even under strong selection, allele transfer into *A. cervicornis* was limited and did not reach fixation within 1,000 years (Fig. 8). Patterns of genome-wide introgression mirrored those at the adaptive locus, with variable niche overlap producing the highest overall ancestry exchange of any simulation, but not exceeding ∼5% on average during the 1,000 year timeframe (Supplementary Fig. 15).

**Figure 8:**
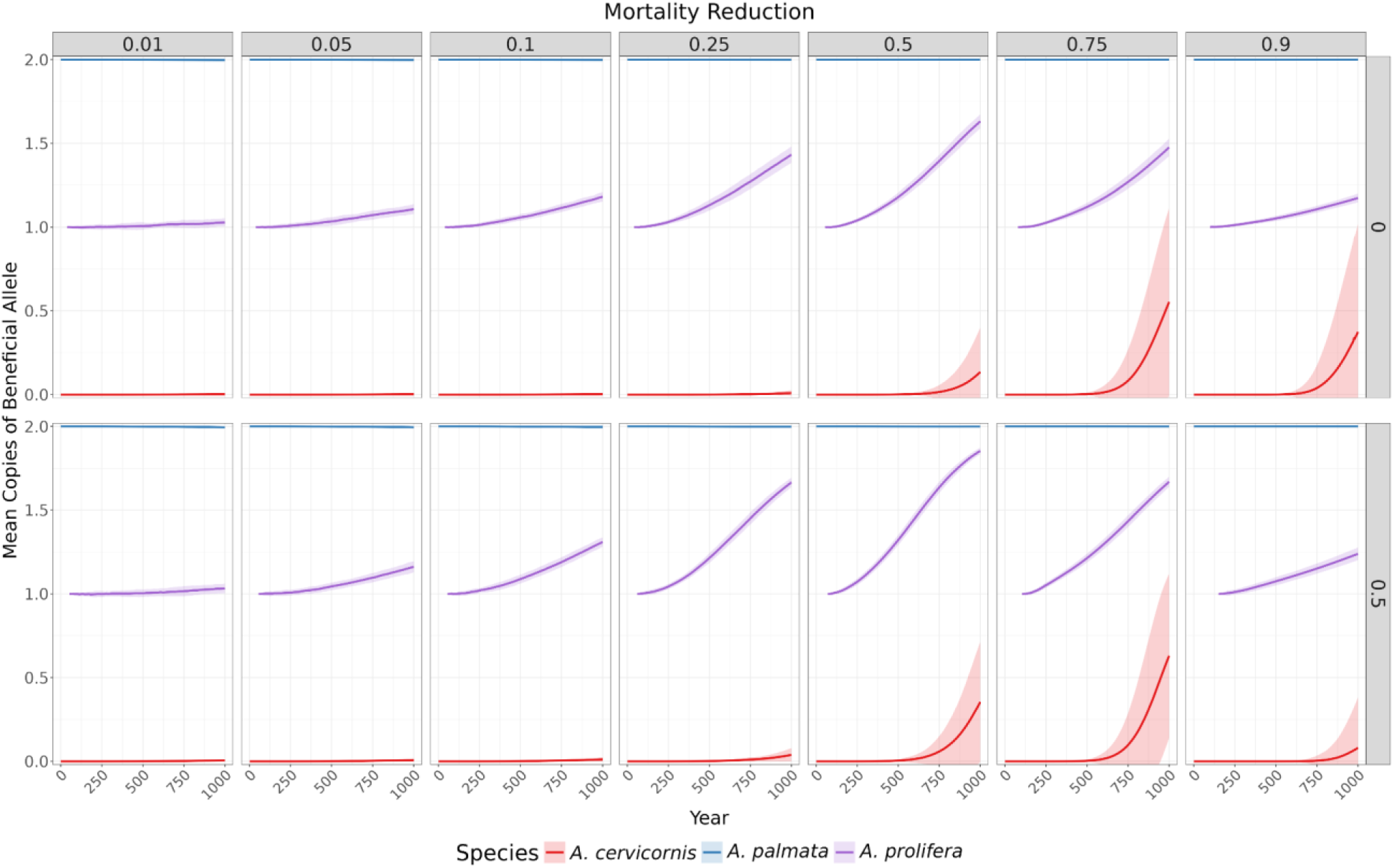
Effect of niche overlaps and single additive beneficial locus on introgression of beneficial locus. The mean number of copies of the additive beneficial allele between replicates (n=50) in the three Acroporids through time, faceted the mortality reduction from carrying the beneficial allele (top) and the niche overlap (right). Blue lines and shading represent *A. palmata*, red lines and shading represent *A. cervicornis*, purple represents *A. prolifera.* Shaded areas show ±2× the standard deviation between replicates (n=50).

Finally, we tested whether allele dominance alternative to the additive model, (dominant vs recessive) influenced adaptive introgression under 50% partially overlapping niches (Fig. 9; Supplementary Figs. 16–17). Strong dominant alleles slowly increased in frequency in the non-adapted species after about 500 years under moderate selection, but not in the hybrids as a single copy already provided full protection. Recessive alleles showed minimal introgression because selection acted only on homozygotes, though the allele became fixed in hybrids in case of strong selection. Genome-wide ancestry followed the same pattern (Supplementary Fig. 16), with the most introgression under dominance, while there was little change in overall species ratios (Supplementary Fig. 17). Introgression was highest under dominance with 50% mortality reduction, but the onset of large-scale gene flow still took approximately 500 years and began to asymptote after 1,000 years when the recipient species was largely outcompeted. Interestingly, our strongest selection strength (90% reduction in mortality) resulted in no genome wide introgression in neither allelic dominance model (Supplementary Fig. 16), again because rapid outcompetition eliminated opportunities for gene flow between parentals (Fig 9; Supplementary Fig. 17). Overall, our results suggest that on short ecological timescales even favorable dominance and partial niche overlap failed to produce meaningful adaptive introgression.

**Figure 9:**
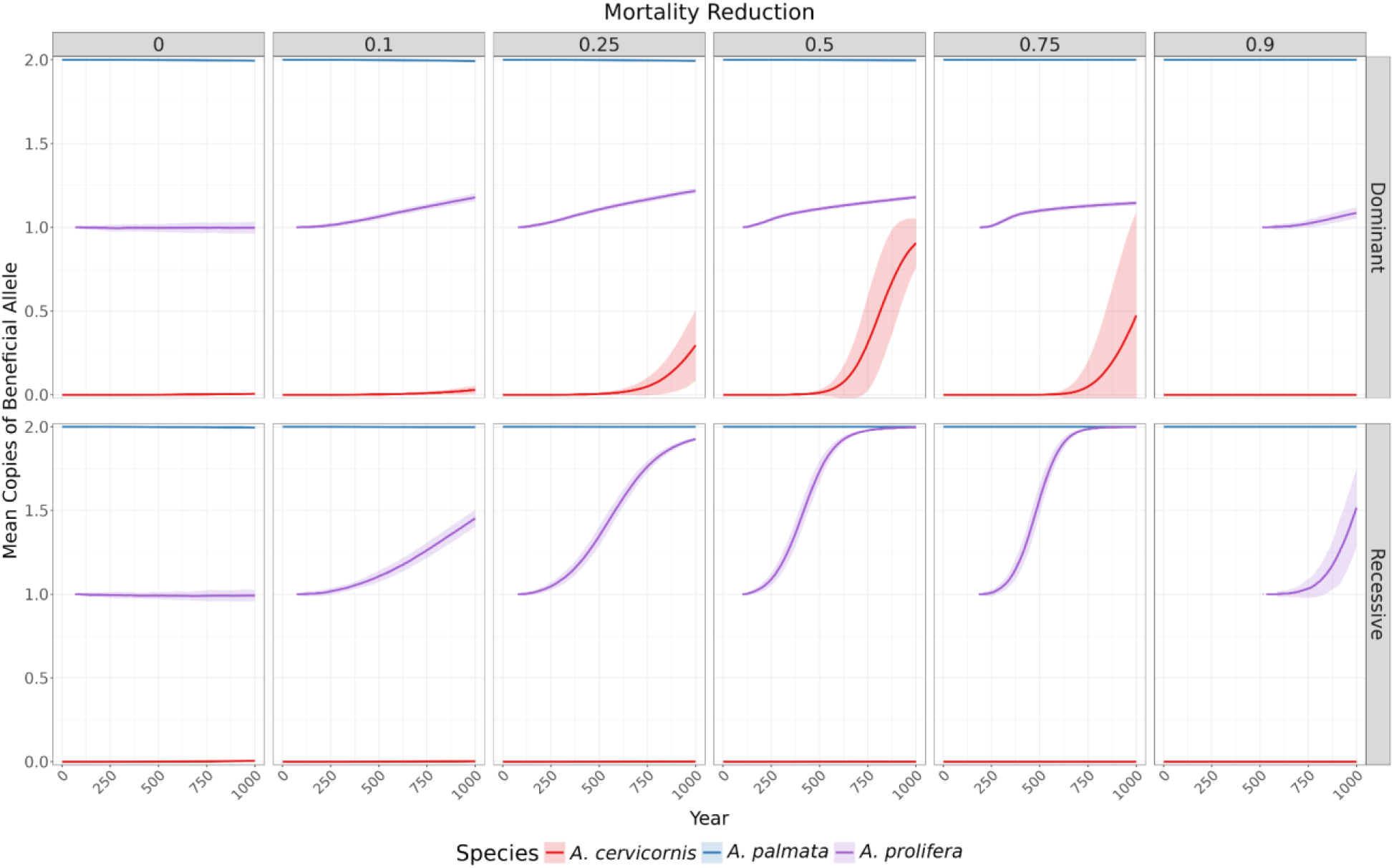
Effect of dominance type of single beneficial locus on introgression of beneficial allele. The mean number of copies of the additive beneficial allele between replicates (n=50) in the three Acroporids through time, faceted by dominance. Blue lines and shading represent *A. palmata*, red lines and shading represent *A. cervicornis*, purple represents *A. prolifera.* Shaded areas show ±2× the standard deviation between replicates (n=50).

## Discussion

The uncertainties surrounding interspecific hybridization in coral restoration are well established in the literature^31,33,35^, and concerns persist about possible ecological or genetic risks associated with introducing hybrids into reef communities^32,35^. To this end, we put forward a two dimensional agent-based simulation model designed to forecast the impact of restoration on species dynamics, hybridization, and rates of introgression within the Caribbean *Acropora* system. We model a system consisting of two species that can hybridize to produce F_1_ offspring, which can subsequently backcross with the parental species. Our models provide little evidence that hybrids become demographically dominant over a 1,000 year timespan, and none in which they completely displace the parental taxa; likewise, we observe little indication of adverse consequences such as genetic swamping.

We find introgression between the two species is generally low, with only a small percentage of heterospecific ancestry being transferred between parental taxa. For ancestry from one parental species to move into the other, hybrids must undergo several consecutive backcrossing events. In our model, we see consistently low heterospecific mating rates, meaning the probability of backcrossing and introgression is low. Further, these results hold in the absence of prezygotic barriers (e.g., mating biases, gametic compatibility, Fig. 2) which all reduce introgression further. The majority of *A. prolifera* individuals in our model are produced by sexual crosses between parentals creating F_1_s, or through asexual reproduction of existing F_1_ hybrid genotypes. As such, the risk of genetic swamping within ecological timescales is predicted to be minimal by our model and suggests that *A. prolifera* may be a viable target for restoration outplanting, as is currently done in very limited contexts (e.g., Laughing Bird Caye in Belize^58^.

This suggests that these biological mechanisms commonly assumed to mitigate introgression^59–61^ may not be the primary determinants of whether genetic swamping may occur in our system; rather, the limiting factor could be the inherently slow, multigenerational process required for heterospecific alleles to rise in frequency. Further, slow accumulation of introgressed ancestry is enforced by demographic priority effects^62,63^ that limit the formation of F_1_ hybrids required for the initiation of heterospecific ancestry transfer. Early numerical dominance of parental taxa is established when parental genotypes are initially outplanted. Because these corals can be long-lived, population turnover and genotype replacement occur slowly. As a result, established colonies retain a persistent demographic advantage.

In our model, clonally generated fragments are able to reproduce with clonemates, in effect increasing the possibility of assortative mating and dampening the impacts of hybridization. However, while selfing in *A. palmata* and *A. cervicornis* possible, it is rare^52,64^. To determine the extent that this dynamic impacts gene flow, we ran simulations where clonal reproduction was allowed (and selfing can potentially occur), and simulations where no clonal reproduction was allowed and selfing was prohibited. In our simulations, we saw that by producing new genetically identical colonies, clonal reproduction allows the existing genotypes to extend their effective genetic “lifespan” within the population— a dynamic that would not occur in obligately sexual species. As implemented, clonal reproduction contributes to buffering populations against genetic swamping.

When clonal reproduction is removed and, as a result, selfing is prohibited, introgression of heterospecific ancestry after 1,000 years increases approximately fivefold relative to the baseline scenarios (maximum of ∼0.1%, Fig 2B., vs maximum of ∼0.5%, Supplementary Fig. 18). Under this scenario we see more hybrids than when clonal reproduction is modeled, however, not enough to exclude parental species (Supplementary Fig. 19). Although this increase in gene flow is substantial, it remains insufficient to cause genetic swamping, demonstrating that other buffering mechanisms alone are sufficient to maintain species integrity even in the absence of clonal reproduction. In long-term simulations (20,000 years), the magnitude of heterospecific ancestry remains similar when clonal reproduction is removed (see Supplementary Fig. 20 and Supplementary Fig. 21). In combination, these results suggest that clonal dynamics do not have a strong impact on gene flow in this system, as the predominant effect observed without clonality is an earlier competitive exclusion of *A. cervicornis* from the reefscape (Supplementary Fig. 22 and Supplementary Fig. 23).

Together, this means that even when conditions favor frequent hybrid formation, the expected genetic outcomes from theory are not realized in the adult population. For instance, broadcast spawning can generate gamete pools that approximate the 50% hybrid proportion expected under random mating (see Supplementary Fig. 3B, column where CSP = 0). Such proportions should rapidly homogenize ancestry in randomly admixing populations^65^. However, these initial mating ratios do not translate into observed population ratios (see Fig. 2C, column where CSP = 0 and where year = 1,000). Instead, priority effects^62,63^ ensure that parental genotypes disproportionately occupy space and reproductive opportunity, reducing the likelihood that hybrids survive, mate, and repeatedly backcross across generations.

Worries about the potential for genetic swamping within this system may be further ameliorated by three additional factors. First, there is an overabundance of *A. prolifera* in our simulations compared to modern real life observations. Some of the most extreme simulations resulted in *A. prolifera* exceeding 50% of colonies (e.g., some simulations from Fig. 3C; Fig. 4; Fig. 7). These relative abundances of *A. prolifera* far exceed historical and modern ecosystem-scale abundances^20,23,66^. This suggests that even with the excess of *A. prolifera* in our simulations (the facilitator of gene flow between the parental taxa), introgression remains low. Second, even when *A. prolifera* abundance is elevated, postzygotic effects such as reduced F_1_ and/or F_2_ viability would even further limit hybrid population growth and reproductive contribution (Fig. 2). Third, to capture occasional long distance clonal dispersal events we simulate a larger clonal reproduction dispersal distance than is likely realistic for most fragments. When we reduce the clonal dispersal range we find even smaller levels of gene flow between parental species (Supplementary Fig. 24), further suggesting that our baseline (Fig. 2B, γ=1) parameters represent a liberal approximation of gene flow.

Although the Caribbean *Acropora* system is among the best-studied coral family with respect to reproductive biology and compatibility, additional laboratory experiments examining conspecific sperm precedence and backcrossing using *A. prolifera* would be highly informative. To date, these processes have been explored in only the Caribbean parental species (e.g., Fogarty et al., 2012^52^). Although it is possible to form F_2_ hybrids from Pacific *Acropora* F_1_s in a lab controlled setting^67^, understanding the genetic breakdown of F_2_ for the Caribbean Acroporids remains unresolved, as there are no published experiments assessing the viability of backcrosses and F_2_s in this system. However, publicly available data from STAGdb^68^ suggests that putative backcrosses are exceedingly rare, with first generation backcrosses representing only 3 of 3,802 genotyped samples and no documented F_2_s (*accessed,* September 2024). This limited evidence suggests that, while F_1_ hybrids are viable, F_2_s may be inviable or possess low fitness and gene flow may be facilitated only by backcrossing, which buffers Dobzhansky-Muller incompatibilities that would otherwise be present (see Edmands 1999^69^ and Yu et al., 2022^70^). By suppressing the number of hybrids that survive to maturity, early-life fitness costs (i.e., Fig. 2E–H) may act in concert with demographic processes to maintain low hybrid frequencies across generations in the wild.

Consistent with the low level of gene flow, the potential for adaptive introgression from one parental species to the other is also limited. Although allele dominance influenced allele dynamics under partial niche overlap—with dominant alleles showing the greatest introgression and recessive alleles showing little—even favorable dominance and strong selection failed to produce meaningful adaptive introgression. Introducing a highly beneficial allele into one parental population gives that species a competitive advantage, allowing it to outcompete the other species. Instead of facilitating gene flow, a highly adaptive allele accelerated demographic dominance and rapidly reduced the potential recipient species abundance and mating opportunities. As a consequence, the recipient species typically did not survive long enough to receive and benefit from the advantageous allele (e.g., Fig. 6). This suggests that competitive dynamics can generate barriers to adaptive introgression within this system, even in the absence of intrinsic reproductive isolation. These findings have direct relevance for genetic-based restoration strategies in corals, where outplanting individuals of one species carrying beneficial variants may enhance one population’s resilience, yet inadvertently suppress the persistence of sympatric taxa before introgression can occur.

Another important consideration for restoration in this sympatric system is how local environment and ecology shape interspecific competition and introgression. In the previously discussed simulations, we assumed fully sympatric outplanting and even parental species outplanting ratio. In addition to this, we model restoration scenarios that vary outplanted species ratios and the degree of niche overlap (Fig. 3), spanning conditions from monotypic to mixed-species plots as observed *in situ* (e.g., Nylander-Asplin et al., 2021^22^). We find that when parental species occupy fully independent non-overlapping niches, both reach carrying capacity within their respective environments (Fig. 3C–D). Ecologically, these fully non-overlapping niches in our simulation would represent parental taxa outplanted across a depth gradient where parental taxa outcompete the other in their local environment (i.e., niche) and the hybrids being equally capable to colonize across niches.

However, in the wild, niches of the parental species often show low to moderate overlap, rather than fully independent non-overlapping niches, and the F_1_ often strongly overlaps with one or both of the parent species (see Nylander-Asplin et al., 2021^22^). In this situation, we see prolonged coexistence of all three taxa, and hybrid lineages were most successful when overlap was substantial, but not complete. Partial niche overlap (where some degree of space competition occurs between parentals) prevents either parental species from fully monopolizing shared space, allowing hybrids to persist and expand.

Importantly, we also find that when one parental species is strongly favored in the initial outplanting ratios and niche overlap is complete (Fig. 3A–B), this numerical advantage often translates into greater long-term success of that species (Fig. 3A). Gametes from the minority taxon are more likely to encounter heterospecific gametes because the majority taxon contributes a disproportionately large number of gametes to the water column. This increases hybrid formation and further accelerates the decline of the minority taxon, as its gametes less frequently result in conspecific reproduction. However, the effects of biased outplanting can be reduced when it coincides with partial niche overlap (Fig. 4). In these cases, early numerical dominance allows the favored species to expand beyond its core niche into shared habitats, temporarily excluding the other species (Supplementary Fig. 10). However, over longer timescales, the initially disadvantaged species persists in its independent niche and eventually re-enters the overlap zone, increasing competition and contact between taxa (Fig. 4). This delayed re-entry promotes hybridization, as both parental species and hybrids co-occur in shared habitats. Hybrids can then spread into both parental niches, reshaping long-term community composition. Notably, these dynamics unfold slowly, becoming apparent only after ∼200 years (Fig. 4). Together, these results show that strongly biased outplanting combined with partial niche overlap can initially favor one species, but ultimately promotes hybrid persistence and introgression across habitats— however, often on timescales longer than those relevant for management.

Although outcomes within the typical duration of our simulations did not differ dramatically across parameter values, this raised the question of whether the system would eventually stabilize—and if so, on what timescale. Extending simulations revealed that while outcomes over the first ∼1,000 years were often similar across a wide range of parameter values, they diverged substantially over longer periods (Supplementary Figure 23). In many cases, relative abundances and community composition continued to shift for 10,000–20,000 years before approaching equilibrium, with long-term states differing markedly from both 1,000-year outcomes and (Fig 2A), more extremely, from those observed over 200-year timescales (Supplementary Fig. 1A). Running the niche overlap and outplanting ratio sims for 6,000 years (or 6× longer than our standard sims), we see that under certain outplanting ratios and niche overlaps, the timing of equilibrium is dependent on the parameter combination (Supplementary Fig. 25). However, this delay and variability in equilibrium is unsurprising, as the models presented here represent a liberal approximation of the *Acropora* system by modelling the secondary contact of two “naive” parent lineages coming into contact for the first time, where in reality the species have coexisted for 2.6-6.6 million years^71,72^.

While we do not place considerable weight on the results of long-term simulations past a thousand years due to the potential for environmental change that cannot be accounted for, long-term simulations do provide useful insight to the past and present of hybridization within the study genus. For instance, long-term simulations suggest that when *A. prolifera* gamete success is equivalent to the parental species (γ=1), *A. prolifera* is able to outcompete the parental species and becomes the only taxon present in the simulated reefscape (Supplementary Figure 23). We know this to not be possible given that all three taxa have existed in sympatry for millions of years^71,72^. Further, *A. prolifera* still remains rare at the ecosystem-level in the present. This suggests that *A. prolifera* gamete success is likely < 0.5 compared to the parental species, or potentially considerably lower, and these simulations provide evidence that, while prezygotic barriers are largely absent between *A. palmata* and *A. cervicornis*^52^, prezygotic barriers such as chromosome missegregation or asymmetric recombination may begin to manifest in *A. prolifera* and limit the generation of F_2_s and backcrosses in nature. Survival reduction of F_2_s (Fig. 2F) did not result in appreciable differences in gene flow or abundance, lending further support that gamete viability in *A. prolifera*, in combination with other parameters such as niche overlap, may be a leading driver of low gene flow in the system. Further experimental work should be performed using standardized crosses to confirm reduced gamete performance in *A. prolifera*.

Finally, we examined how project size, measured by the number of outplanted corals, affects restoration outcomes^73^. Larger outplanting efforts showed much lower variability in final population size (Fig. 5), reflecting reduced sensitivity to stochastic demographic and ecological processes that strongly affect small founding populations^74,75^. While mean species composition and introgression were similar across outplant sizes, replicate-to-replicate variability depended strongly on initial population size. Small outplantings often diverged in long-term trajectories, whereas planting several hundred individuals produced consistent outcomes across replicates (Fig. 5A–B), indicating a practical threshold for predictable restoration. Notably, these stochastic effects became apparent only after ∼200–250 years, well beyond typical monitoring timescales. Although our closed-system simulations likely overestimate stochasticity compared to well-connected reefs, the key result—that larger outplant sizes reduce stochastic divergence and improve predictability—should hold, especially for isolated or degraded reefs.

While our simulations offer a controlled framework for exploring long-term outcomes, field conditions are inevitably more complex. Future work could bridge this gap by collaborating with restoration practitioners to conduct small-scale outplanting trials that test whether simulation predictions hold under *in situ* conditions. This would require outplanting, monitoring and sequencing to determine ancestry patterns within the population. In practice, *in situ* confirmation may be challenging due to ocean currents driving larval export and import(e.g., Galindo et al., 2006^76^), as well as the well-documented chronic recruitment failure of broadcast-spawning stony corals in the Caribbean^77^. Additionally, our models simulate “set it and forget it” strategies, where restoration takes the form of an initial strategic outplanting, followed by no intervention. Our results do not preclude the possibility of more active management and breeding strategies (e.g., assisted gene flow between populations^78,79^ or adaptive gene flow between species^80,81^ through targeted crossing) producing meaningful results that increase the resilience of any and all members of the genus.

Although our study focuses on Caribbean *Acropora*, our modeling framework is broadly applicable. Many coral systems exhibit histories of hybridization and cryptic speciation^32,38,59^. For example, the framework could be applied to other simple but threatened systems such as *Orbicella*, an increasingly important target of reef restoration following the functional extinction of *Acropora* in Florida^59,82–85^. As in *Acropora*, gene flow among the three *Orbicella* species is thought to occur based on laboratory compatibility experiments, although confirmed hybrids have not been identified in the wild, potentially due to difficulties in morphological identification^59,85^. Evolutionary simulations in SLiM^47^, combined with targeted field experiments, could help clarify patterns of gene flow and their implications for restoration in *Orbicella*.

More broadly, this work highlights the value of simulation-based approaches in generating long-term evolutionary and ecological insights that are otherwise inaccessible through field observation alone. By continuing to integrate empirical data into dynamic models, restoration science can better anticipate the consequences of human interventions and refine strategies to support coral resilience in a rapidly changing ocean.

## Methods

### Model overview

Our simulation model is written in the Eidos language^86^, which is designed for the SLiM simulation framework (v4.0). SLiM is a software that provides users the ability to code detailed population genetics simulations^47^. Our forward-in-time agent-based simulation consists of overlapping generations and is designed to mimic coral biology as closely as possible. We model the impact of different restoration regimes on each *A. cervicornis, A. palmata*, and *A. prolifera* from a genomic perspective, while incorporating sexual and asexual reproduction, mortality, settlement, and competition. Each individual coral has a location within a coordinate plane representing reef substrate. The size of the coordinate plane is based on a simplified rectangular approximation of a large, hypothetical Caribbean restoration site of approximately 13km by 9km. All simulations referenced in the main text figures were run in replicates of 50 except the “variable outplant size” simulations which were run in replicates of 150 due to the increased variance under small outplanting projects. Unless otherwise specified, simulations were conducted using a baseline parameter set (niche overlap = 1; γ = 1; *A. palmata : A. cervicornis* outplanting ratio = 50:50; CSP values of *A. palmata* = 0.333 and *A. cervicornis* = 0.03; no adaptive loci and no increased bleaching mortality except in simulations explicitly designed to test those effects—see parameter descriptions in sections below). Individual restoration parameters were then varied one at a time while holding all other parameters at their baseline values to isolate their independent effects on model outcomes.

### Mechanisms of genetic isolation between species, reproduction, and dispersal

To evaluate how variation in *A. prolifera* gamete success influences introgression rates and species dynamics, we incorporate a gamete success parameter (γ) into the model. This parameter modulates the relative success of hybrid sperm and eggs and is motivated by the assumption that *A. prolifera* gametes may experience reduced viability (a prezygotic form of hybrid breakdown) compared to parental gametes due to asymmetrical recombination in hybrids^53–55^. We examined γ values ranging from 0 to 1 (0, 0.25, 0.5, 0.75, 1.0). In the model, γ = 1 indicates that hybrid gametes have no fitness disadvantage relative to parental gametes. In contrast, γ = 0 represents complete gametic failure of *A. prolifera*, such that hybrids cannot produce viable offspring (i.e., neither backcrosses nor F_2_ individuals are possible, as both require functional hybrid gametes).

Additionally, we incorporate conspecific sperm precedence (CSP) into the reproduction function. This models the capacity of eggs to preferentially recognize and fertilize with conspecific sperm rather than heterospecific sperm. The implementation is informed by Fogarty et al., (2012) which conducted mixed sperm trials with *A. palmata* and *A. cervicornis* eggs^52^. While our baseline CSP values for *A. palmata* (CSPpal = 0.333), and *A. cervicornis* (CSPcer = 0.03) are based on Fogarty et al., (2012)^52^, we also test variable symmetrical values of both CSPpal and CSPcer (0, 0.25, 0.5, 0.75, 1.0), to assess the impact of conspecific sperm precedence on the level of introgression.

Corals in our model produce offspring in two ways: sexual reproduction via broadcast spawning, and asexual reproduction via fragmentation. In studies across the Caribbean, the ratio of sexually and asexually derived colonies is roughly 1:1^48,87,88^, meaning that clonal reproduction and fragmentation are equally successful on the reefscape. However, in degraded or bottlenecked populations (e.g., Florida or Guadeloupe), the rate of asexual reproduction was reported to be 79× higher than sexual reproduction in *A. cervicornis*^88^ and 19× higher in *A. palmata*^87^. Because the site we model here is meant to represent a site in need of restoration, we assign clonal reproduction a modestly higher success rate (2×) than that of sexual reproduction, although we also include scenarios with no clonal reproduction to understand the impact on hybridization dynamics (Supplementary Figs. 18–19).

In the case of sexual reproduction, offspring genomes are generated via recombination of the parental genomes (see *Genome Modeling* section for more details). In the case of fragmentation reproduction, the offspring genome is a clone of the individual from which it originally broke away. Because reproductive capacity depends on both age and colony size, we represent a size-dependent reproductive threshold in the model using age thresholds, such that reproductive age serves as an operational proxy for the minimum colony size and age required for sexual reproduction and fragmentation.

We assume that corals must be at least 8 years old to reproduce sexually based on a long-term study from Mexico that followed outplanted *A. palmata* sexual recruits^89^, and at least 2 years old to reproduce asexually via fragmentation. Consequently, younger corals are unable to reproduce and for these corals to reach reproductive maturity, they must survive across successive years (see *Mortality* section for more details). To our knowledge, no study exists for *A. cervicornis* that estimates the time to reproduction starting from a sexually-generated recruit, so we assume that the time to sexual reproduction is similar for the two species.

New asexual fragments produced within a given year are assigned an initial age of 2 years to account for accelerated physical and sexual development, while sexually produced larvae are assigned an initial age of 0 years. This means that asexual fragments are capable of reproducing sooner than larval recruits—being able to reproduce after ∼6 years based conservatively on Williams et al., (2023)^90^. For coral fragments, we assume the lost biomass is equivalent to the growth required to reach the size of a 2-year-old coral. We therefore reduce the effective age of the parent colony by 2 years so that its post-fragmentation size corresponds to that of a colony two years younger, also reflecting the short-term energetic cost of tissue loss and regrowth^91^. Lastly, we assume that corals which have been outplanted at the beginning of the simulation start at 4 years of age, meaning it takes at least 4 years following outplanting for sexual offspring to be produced^91^.

Each year, to facilitate sexual reproduction, each reproductively aged individual spawns by releasing male and female gametes into the water column. Distances between reproductive colonies greatly influence the number of offspring produced for broadcast spawning marine organisms^61^. Gametes generally have small dispersal distances because they are short-lived and quickly lose viability in the water column^92^.

To calculate radius of gamete dispersal, *m*, we retrieved average surface ocean current direction and strength data from HYCOM^93^, taken over the duration of the coral spawning period in Belize (August 11, 2022–August 23, 2022). Over the spawning period, the average velocity from HYCOM was 0.0567 meters/second, respectively. This value, combined with the maximum duration gametes are viable (∼3hrs^92^), allowed us to calculate the maximum distance gametes could theoretically travel (∼0.6km), which became our maximum sexual interaction distance (*m* = 0.6km).

To simulate fertilization, we model the probability of egg-sperm fertilization (i.e., accounting for gamete-level prezygotic selection) based on a weighted sample where sperm are selected for by eggs. Specifically, sperm are sampled for fertilizing eggs at weights described by equations 1.0–3.2 below. These equations include reproductive isolation parameters CSP (which is denoted per species with subscript species abbreviation) and γ. The equations also consider the local number of individuals of each species within *m*. We refer to this local number of reproductively aged individuals using the parameter *ρ* (denoted per species with subscript species abbreviation).

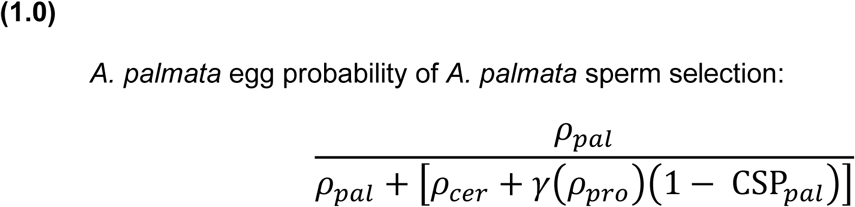

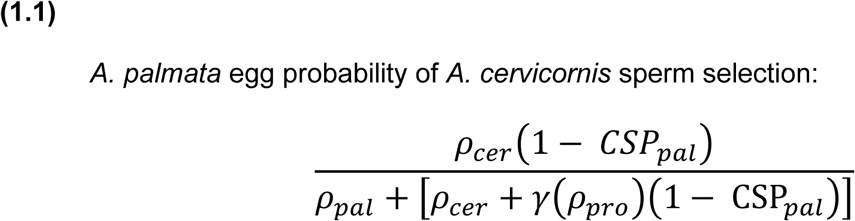

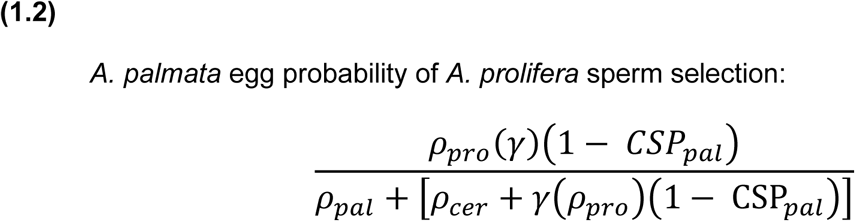

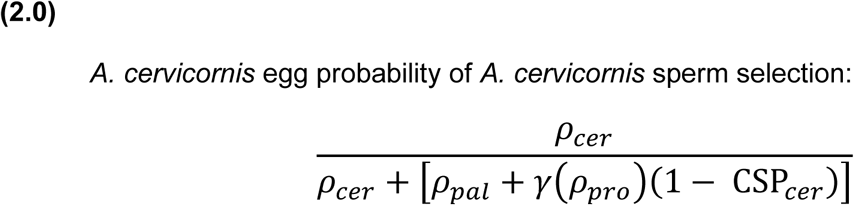

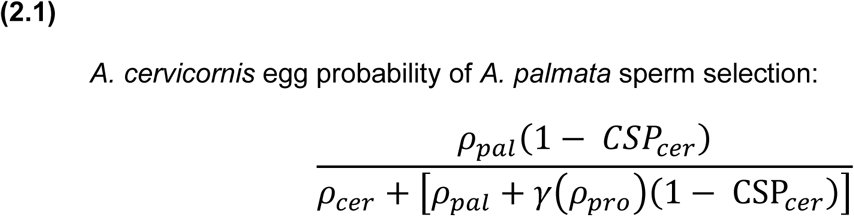

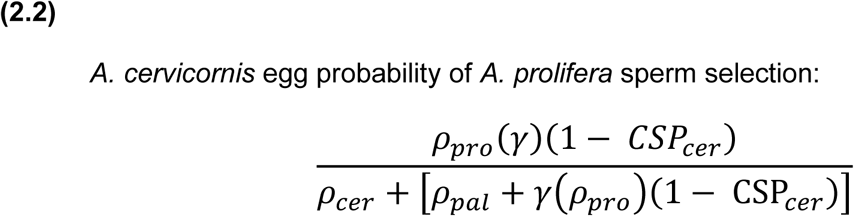

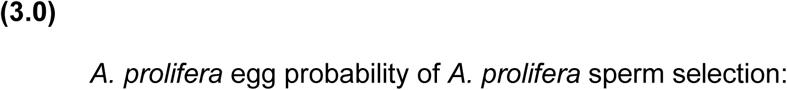

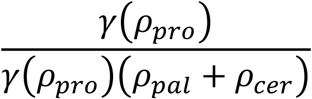

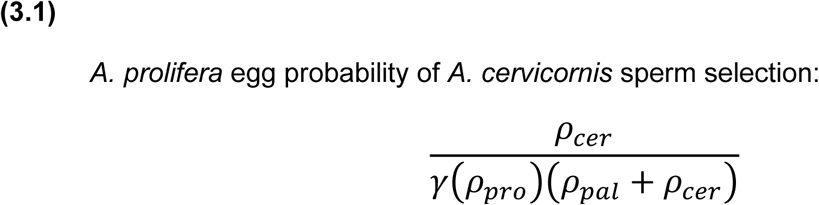

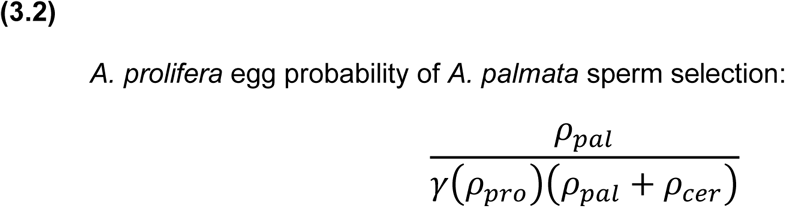

Unlike gametes, larvae can remain in the water column for extended periods and disperse over greater distances^25,94^. *Acropora* larvae have been observed to survive for approximately 4– 5 days^25^, and studies have consequently estimated dispersal distances on the order of several kilometers(Baums, Hughes, et al., 2005; Baums, Miller, et al., 2005; Drury et al., 2018; Hemond & Vollmer, 2010), although much longer larval duration periods are possible, but rare(e.g., Graham et al., 2008^98^). Dispersal distances for fragments and larvae are modeled using Rayleigh distributions, representing how far each agent can travel within the water column. This Rayleigh formulation arises from assuming that dispersal occurs isotropically in two dimensions, with horizontal displacements in the X and Y directions drawn independently from normal distributions with mean zero and standard deviation σ.

In addition to sexual reproduction, branching corals such as Acroporids also reproduce asexually through fragmentation, whereby fragment agents break off from a parent coral agent and can settle to generate a new colony. Studies of clonal diversity in *A. palmata* suggest that ramets of a genet can be as far as ∼100m from one another^87^. Although not specifically documented for Caribbean *Acropora*, rolling coralliths have been documented in the Caribbean and elsewhere and have the potential for much longer dispersal distances^99,100^. For all dispersal events, an agent’s post-dispersal location is calculated by adding the sampled X and Y displacement values to its initial (X, Y) coordinates.

For the reasons stated above, standard deviations of the dispersal distribution (σ) are agent-type specific and parameterized such that fragment dispersal occurs at a spatial scale 20-fold smaller than larval dispersal, capturing spatial differences among reproductive modes. These values (0.5km and 10km for fragments and larvae, respectively) represent an effective dispersal kernel that integrates both background dispersal and occasional long-distance recruitment events (e.g., storm- or swell-driven transport), thereby reflecting dispersal under variable hydrodynamic conditions across the 1,000 year simulation. We also reduced the fragment dispersal scale by an order of magnitude (i.e., 10× smaller than the baseline) and found that key model outcomes were qualitatively unchanged, with these results including even lower gene flow indicating robustness of the findings (Supplementary Fig. 22).

### Mortality

After gamete release and fertilization, colonies can produce larvae. We do not explicitly simulate gamete number (eggs or sperm) or fertilization dynamics at the level of individual gametes. Instead, reproduction is modeled in a coarse-grained manner: each reproductively mature colony makes a recruitment attempt per year. If compatible sperm is available from nearby mature neighbors, the colony produces a larval agent representing the outcome of its spawning event (i.e., at least one successful fertilization and many mortality events among many gametes). The model’s single larva per colony per year should not be interpreted as a single egg; it is a computational proxy for a spawning event producing many gametes, summarized to a potential recruit. However, a single colony may contribute genetically to multiple larvae within the same year as a sperm donor because it can be sampled repeatedly by different neighboring egg parents. Immediately after larval production, we apply strong early mortality by allowing larvae to survive with probability 1%, corresponding to 99% early mortality, capturing high post-fertilization attrition prior to settlement. Surviving larvae then disperse, must land on suitable habitat, and must pass a local density settlement filter (see *Settling, Competition and Niche* section). This is consistent with studies that show high mortality in larvae post settlement^101,102^.

After this initial mortality step, the survival of the remaining offspring (both fragments and larvae) is dependent on there being suitable habitat for offspring to settle. If larvae move “off the reef” (i.e., into the open ocean or washes up on land) they die. This is represented on the model landscape by an X, Y coordinate that is not within the bounds of our hypothetical reef. If larvae do end up on reef substrate, but there is no available space to settle (because the local population density where the larvae/fragment settled is too high) the offspring do not settle effectively, and also die (see *Settling, Competition and Niche* section). These forces within our simulation allow us to maintain the local population density.

Finally, if a larva or fragment successfully settles and survives to adulthood, it is subject to background adult mortality. Because of the way that sexual reproduction is modeled, wherein only one recruit is generated per pairing of colonies, we were required to reduce the magnitude of annual probability of mortality from 10% (see Riegl and Purkis 2015^103^) to 0.1%, otherwise simulated populations were prone to collapse (total loss of all species). In non-bleaching years, each adult colony faces a fixed 0.1% annual probability of death (99.9% survival), reflecting long colony lifespans and low demographic turnover. Under these rates, expected annual adult losses are lower than potential recruitment from sexual reproduction, with additional input from clonal propagation. Consequently, in the absence of disturbance, population dynamics are regulated primarily by space limitation and post-settlement density dependence rather than reproductive output.

In the adaptive introgression “bleaching simulations”, mortality dynamics are modified further to reflect environmental stress. During bleaching years, adult mortality is assumed to double (see bleaching mortality statistics from Harriott (1985)^104^ and Riegl & Purkis (2015)^103^) to 0.2%. However, individuals carrying a hypothetical beneficial resistance allele experience reduced bleaching-related mortality. We simulated the spread of such an allele under varying selective strengths, where carrying the allele reduced the probability of adult mortality during bleaching events by 1%, 5%, 10%, 25%, 50%, 75%, or 90%. Individuals homozygous for the allele received the full mortality reduction whereas heterozygotes received half of that benefit. We also tested alternative dominance scenarios (dominant and recessive) to evaluate how allele dominance influenced adaptive introgression.

The frequency of large-scale bleaching events follows estimates from Hughes et al. (2018), who reported bleaching occurring approximately once every 5.9 years^105^. Accordingly, the baseline bleaching probability was set at 16.7% per year, and increased by 4% annually, as projected by Hughes et al. (2018)^105^, until reaching a maximum of 100%—representing bleaching conditions occurring every year.

To examine postzygotic isolation via reduced hybrid viability, we implemented two simulation scenarios resulting in increased hybrid mortality. In both cases, we retained our baseline larval survival of 1% for parental genotypes. In the first scenario (reduced F_2_ success), F_1_ hybrids and backcrosses exhibited baseline survival equivalent to parental genotypes, whereas F_2_ hybrids experienced increased mortality compared to parentals, which captures delayed postzygotic isolation arising from recombination-dependent genetic incompatibilities (see *Genome modeling*). F_2_ larval success was reduced multiplicatively relative to the baseline by factors of 5, 10, 100, or 1,000 (i.e., 1% divided by 5, 10, 100, or 1,000), yielding survival probabilities of 0.2%, 0.1%, 0.01%, and 0.001%, respectively. In the second scenario (general hybrid inferiority)—which represents immediate and persistent postzygotic isolation—all hybrid classes (F_1_, F_2_, and backcrosses) experienced reduced larval survival. We decreased hybrid viability stepwise by halving the baseline value (1%) yielding 0.5%, 0.25%, 0.125%, and 0.0625%.

### Settlement, Competition, and Niches

Local density maxima (carrying capacity, *K*) were parameterized using spatially explicit genet data based on work from Drury et al., who mapped the distribution of unique *A. cervicornis* genotypes across surveyed reefs within defined spatial plots^48^, allowing estimation of the number of unique genets per unit area. In the model, each individual represents a single coral unit in SLiM^47^, but translating this to field density is non-trivial because coral colonies vary substantially in size and a single colony can occupy variable amounts of space. To anchor model density to empirical observations, we therefore used the density of unique genets as a proxy for how many distinct coral units can co-occur within a given area. We converted the maximum observed density of unique genotypes in these mapped areas to individuals per km^2^, yielding *K* = 262 individuals/km^2^. Because of the lack of similar, spatially explicit data in *A. palmata* and *A. prolifera*, we assume the estimated density of *A. cervicornis* serve as a realistic proxy for the closely related

### A. palmata and A. prolifera

Settlement is density-dependent and constrained by local space availability. To represent local crowding, we define a circular interaction distance around each individual with radius *s,* which we assume to be 1km. We calculate the area of this interaction zone as 𝜋*s*². Multiplying this area by *K* gives μ, or the expected number of corals that can occupy that circular neighborhood at carrying capacity. Thus, μ represents the maximum number of individuals within the competition circle, and is used to determine whether sufficient reef space is available for larval settlement. If the number of individuals within the interaction radius reaches μ, no additional larvae can settle in that area. This enforces an upper limit on the number of individuals within each competition neighborhood, consistent with *K*.

Importantly, competitive interactions are further structured by ecological niche differentiation. Evidence indicates that *A. palmata* and *A. cervicornis* occupy slightly different positions along a depth gradient. Although the two species are largely sympatric, *A. palmata* preferentially occupies the shallowest depths within the shared range^49–51^, effectively reducing direct competition for space and influencing local settlement dynamics.

As such, we model varying degrees of niche overlap between planted corals. At one extreme, we simulate completely independent *A. palmata* and *A. cervicornis* niches, in which parental species compete only with conspecifics for reef habitat, and the only interspecific competitor is *A. prolifera*. At the other extreme, we model fully overlapping niches (100% overlap), in which parental species are fully sympatric, and compete directly for space. Intermediate niche overlaps (25%, 50%, and 75%) are also explored to assess how partial niche independence influences population dynamics.

To represent niche differentiation along depth within our simulated two-dimensional reef plane, we incorporate niche overlap as a modifier of local competitive interactions rather than as an explicit third spatial dimension. Local interactions are determined within the same circular interaction distance (radius *s*) used for settlement—within each interaction circle, we calculate a realized local abundance, denoted by η, which represents the effective number of individuals contributing to competition for a given species. Specifically, conspecific individuals within the interaction radius contribute fully to η, whereas heterospecific individuals contribute proportionally according to the niche overlap parameter, λ. Thus, η is the weighted sum of individuals within the interaction circle, where λ determines the strength of interspecific competition. When λ = 1, niches overlap completely and heterospecifics contribute equally to competition, such that all individuals within the interaction radius count fully toward the shared carrying capacity. When λ = 0, niches are fully independent and the heterospecific parental species does not contribute to each other’s effective local abundance.

During settlement, larvae of a given species are permitted to settle only if their species-specific realized local abundance, η, is less than the maximum allowable local abundance, μ, within that interaction circle. If η ≥ μ, settlement fails and the larva dies. This implementation modifies competitive effects without introducing an explicit third spatial dimension and remains consistent with the local carrying capacity framework described above.

### Genome modeling

Each coral has a diploid genome that is 267Mb, or the total size of the 14 pseudochromosome scaffolds assembled in an initial pre-release version of the *A. palmata* genome (now published in Locatelli et al., 2024^106^). Genomes have per-chromosome recombination rates from the initial pre-release version of the *A. palmata* recombination map. The pre-release map differs slightly from the published version (see simulation scripts, *Code Availability Statement*), but all simulation results are analyzed at the genome-wide scale and are not impacted by slight deviations at the chromosome-level. The genome-wide and chromosome-specific recombination rates for *A. cervicornis* and *A. palmata* were found to be similar^106^, and were thus given the same values in simulations. Due to the lack of recombination data in *A. prolifera*, we assumed that recombination rates in the hybrid would be similar to the parentals, although this may not account for missegregation that can occur in divergent sets of chromosomes, as in F_1_ hybrids (e.g., Bozdag et al., (2021)^107^; Rogers et al., (2018)^108^). To capture possible missegregation or general reduced hybrid viability we test separate scenarios in which hybrid gamete success is reduced (see *Mortality* section).

### Ancestry estimation

Ancestry proportions are determined by neutral markers fixed across the entire genome of all *A. palmata* individuals upon initialization of the simulation (i.e., when individuals are outplanted). A single neutral marker mutation is placed every megabase along the 267Mb genome. The proportion of heterospecific ancestry in each individual is calculated in subsequent years by counting the number of markers present in each individual, and dividing by the total number of possible marker locations. If an individual has all of the possible markers, then it has 100% *A. palmata* ancestry. Similarly, if an individual has none of the possible markers, then it has 100% *A. cervicornis* ancestry. Upon initialization all corals are considered to be either 100% pure *A. palmata* or 100% pure *A. cervicornis.* For simulations modeling a single beneficial mutation, we initialize the allele as fixed at a single locus in *A. palmata* and track its frequency over time.

### Species Determination

To determine the species identity of each individual, we use a set of species markers that are distinct from those used for ancestry estimation (described above). This approach is analogous to the STAGdb species identification SNP panel developed for Caribbean *Acropora* corals, which are used to distinguish the species and hybrids in wild-sampled individuals^68^. In our model, species identity is determined by the number of these species markers an individual possesses. Ninety speciation loci were generated, with one every 3Mb on the simulated genome. To be considered as one of the parental species, an individual must have 90% percent of the parental alleles present in its genome (i.e., 162 of the possible 180 alleles must be *A. palmata* alleles for an individual to be considered a parental *A. palmata* and not a hybrid.)

## Author Contributions

T.M.L., C.N.H., N.S.L., and C.D.H. conceptualized and designed the study, T.M.L. developed and analyzed the agent-based simulations, T.M.L. and C.D.H. prepared the manuscript, T.M.L., C.N.H., N.S.L., and C.D.H. revised and edited the manuscript, C.D.H. supervised the study. All authors read and approved the final manuscript.

## Code Availability Statement

The Edios code for the SLiM agent-based model used in this study, a file explaining the code for the SLiM agent-based model—the model, the R data analysis scripts, can be found in the following repository: https://github.com/TroyLaPolice/The_Digital_Reef-Insights_from_agent_based_modeling

