## Supplementary Information for "Simulations reveal hybridization in Caribbean *Acropora* restoration poses low risk of genetic swamping but limited potential for adaptive introgression"

### Table of Contents

#### Supplementary Figures:

*Supplementary Figure 1: Effect of gametic incompatibility on introgression and species dynamics within first 200 years*

*Supplementary Figure 2: Parentage proportions of crosses under different values of hybrid gamete success*

*Supplementary Figure 3: Parentage proportions of crosses under different values of conspecific sperm precedence*

*Supplementary Figure 4: Effect of F2 success reduction and post-zygotic isolation on introgression and species dynamics within first 200 years*

*Supplementary Figure 5: Parentage proportions of crosses under different amounts of F2 success reduction*

*Supplementary Figure 6: Parentage proportions of crosses under different amounts of A. prolifera post-zygotic success*

*Supplementary Figure 7: Effect of initial species ratios and niche overlap on introgression and species dynamics within first 200 years*

*Supplementary Figure 8: Parentage proportions of crosses under different initial species ratios*

*Supplementary Figure 9: Parentage proportions of crosses under different niche overlap values*

*Supplementary Figure 10: Effect of the interaction between initial outplanting ratio and niche overlap on species dynamics within first 200 years*

*Supplementary Figure 11: Effect of outplant project size on restoration outcomes within first 200 years*

*Supplementary Figure 12: Parentage proportions of crosses under different numbers of initially outplanted individuals*

*Supplementary Figure 13: Effect of single beneficial loci initially fixed in A. palmata on introgression and species dynamics within first 200 years*

*Supplementary Figure 14: Parentage proportions of crosses under different values of mortality reduction under an additive mutation model*

*Supplementary Figure 15: Effect of niche overlaps and single additive beneficial locus initially fixed in A. palmata on heterospecific ancestry transfer*

*Supplementary Figure 16: Effect of mutation dominance and single additive beneficial locus initially fixed in A. palmata on heterospecific ancestry transfer*

*Supplementary Figure 17: Effect of mutation dominance and single additive beneficial locus initially fixed in A. palmata on species dynamics*

*Supplementary Figure 18: Effect of hybrid gamete success (γ) on introgression without clonal reproduction*

*Supplementary Figure 19: Effect of hybrid gamete success (γ) on species dynamics without clonal reproduction*

*Supplementary Figure 20: Effect of hybrid gamete success (γ) on long term introgression (20,000 years) without clonal reproduction*

*Supplementary Figure 21: Effect of hybrid gamete success (γ) on long term introgression (20,000 years)*

*Supplementary Figure 22: Effect of hybrid gamete success (γ) on long term species dynamics (20,000 years) without clonal reproduction*

*Supplementary Figure 23: Effect of hybrid gamete success (γ) on long term species dynamics (20,000 years)*

*Supplementary Figure 24: Effect of smaller clonal dispersal distance on introgression levels*

*Supplementary Figure 25: Effect of the interaction between initial outplanting ratio and niche overlap on long term species dynamics (6,000 years)*

### Supplementary Figures:

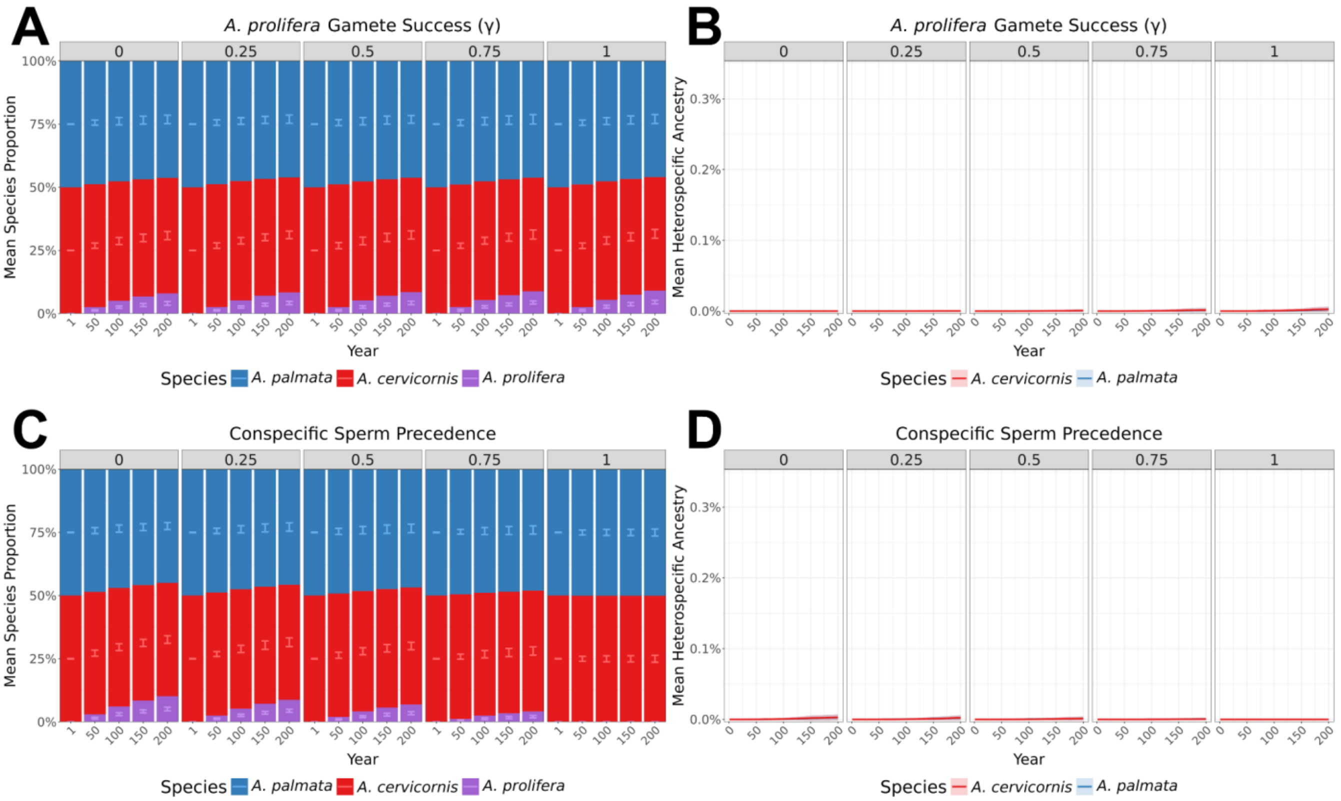


***Supplementary Figure 1:*** **Effect of gametic incompatibility on introgression and species dynamics within first 200 years**

Compared to Main Text Fig. 2A–D, this plot is the same data but instead focusing on the first 200 years of the simulation (i.e., a shorter timespan than Main Text Fig. 2A–D). Species ratios (A, C) and mean heterospecific ancestry (B, D) through time faceted by hybrid gamete success (γ) (A, B), and conspecific sperm precedence (CSP) (C, D). For species-ratio panels (A, C), the x-axis shows simulation years and the y-axis the mean species ratio across replicates (n=50). Stacked bars display the mean proportion of each taxon, with error bars indicating ±2× standard deviation between replicates. Colors: blue = *A. palmata*, red = *A. cervicornis*, purple = hybrid *A. prolifera*. For heterospecific-ancestry panels (B, D), solid lines represent mean heterospecific ancestry across replicates (n=50) and shaded areas show ±2× the standard deviation between replicates. Colors: blue = *A. palmata*, red = *A. cervicornis*.

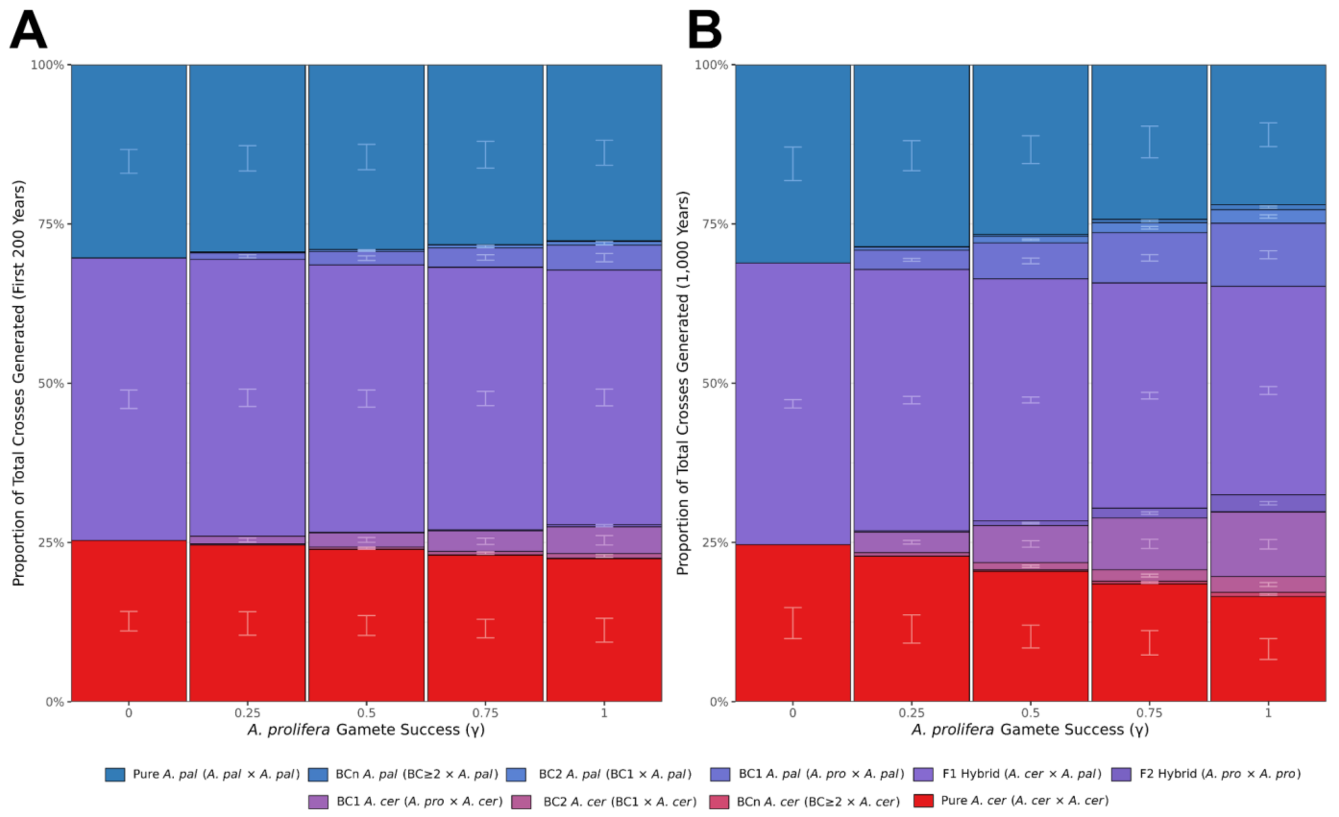


***Supplementary Figure 2:*** **Parentage proportions of crosses under different values of hybrid gamete success**

Unlike the standard species dynamic ratio plots, the x-axis of this figure represents the hybrid gamete success (γ) rather than the simulation year (γ = 0 hybrid gametes are not viable, γ = 1 hybrid gamete viability equal to parentals). Additionally, compared to standard species dynamic plots, instead of there being three categories (with blue bars representing *A. palmata*, red bars representing *A. cervicornis*, and purple bars representing *A. prolifera*), these stacked bars break out the hybrid *A. prolifera* bar into four different bars representing $F_1$s, $F_2$s, $BC_1$s and $BC_2$s. The parental bars are broken into two separate categories, $BC_3$+ and pure parentals with no history of backcrossing. The stacked bars represent a mean of the total proportion of the respective crosses across 50 replicates. The error bars represent ±2× the standard deviation between replicates. (A) Shows the total parentage proportions generated in the first 200 years of the simulation. (B) Shows the total parentage proportions generated over all 1,000 years of the simulation.

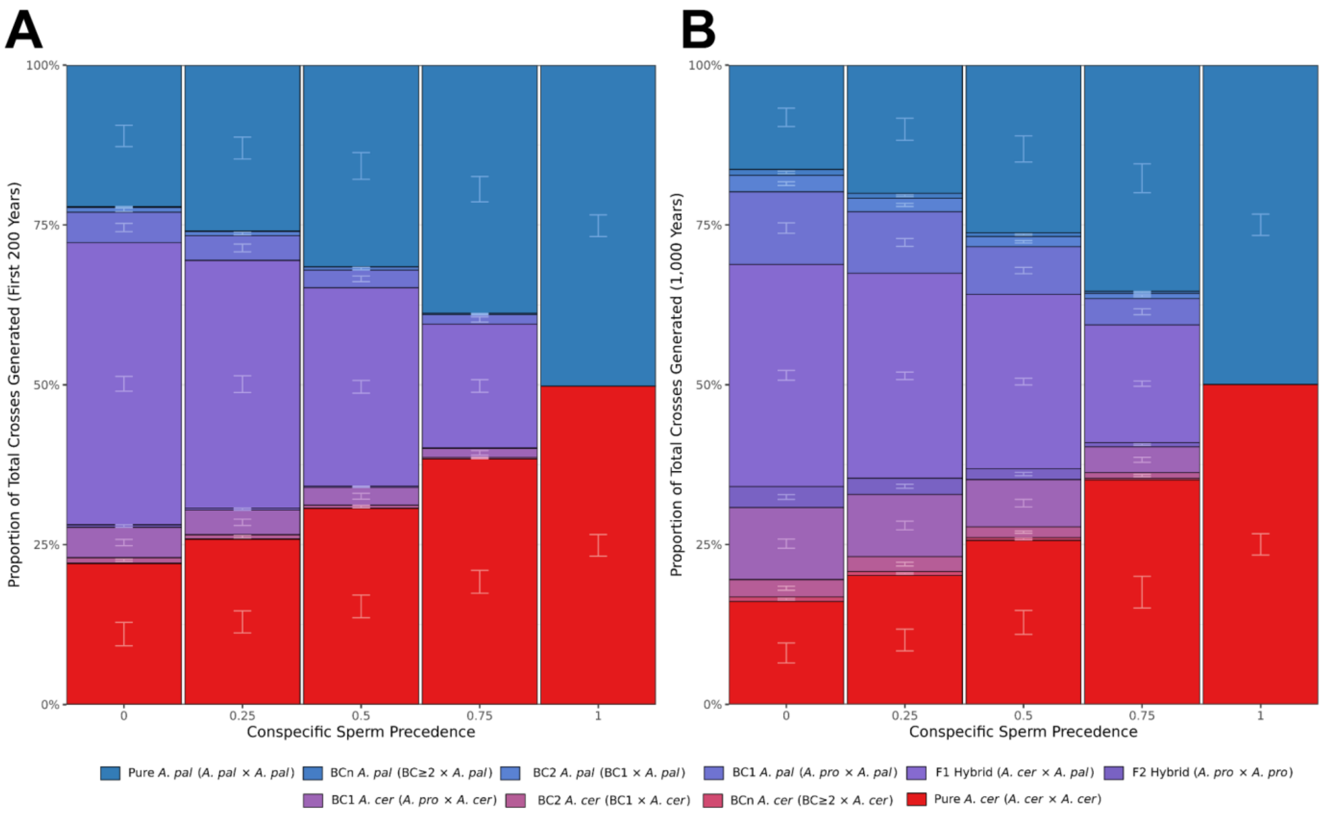


***Supplementary Figure 3:*** **Parentage proportions of crosses under different values of conspecific sperm precedence**

Unlike the standard species dynamic ratio plots, the x-axis of this figure represents the conspecific sperm precedence (CSP) value rather than the simulation year. CSP = 0 represents no sperm preference whereas CSP = 1 represents only conspecific mating. Additionally, compared to standard species dynamic plots, instead of there being three categories (with blue bars representing *A. palmata*, red bars representing *A. cervicornis*, and purple bars representing *A. prolifera*), these stacked bars break out the hybrid *A. prolifera* bar into four different bars representing $F_1$s, $F_2$s, $BC_1$s and $BC_2$s. The parental bars are broken into two separate categories, $BC_3$+ and pure parentals with no history of backcrossing. The stacked bars represent a mean of the total proportion of the respective crosses across 50 replicates. The error bars represent ±2× the standard deviation between replicates. (A) Shows the total parentage proportions generated in the first 200 years of the simulation. (B) Shows the total parentage proportions generated over all 1,000 years of the simulation.

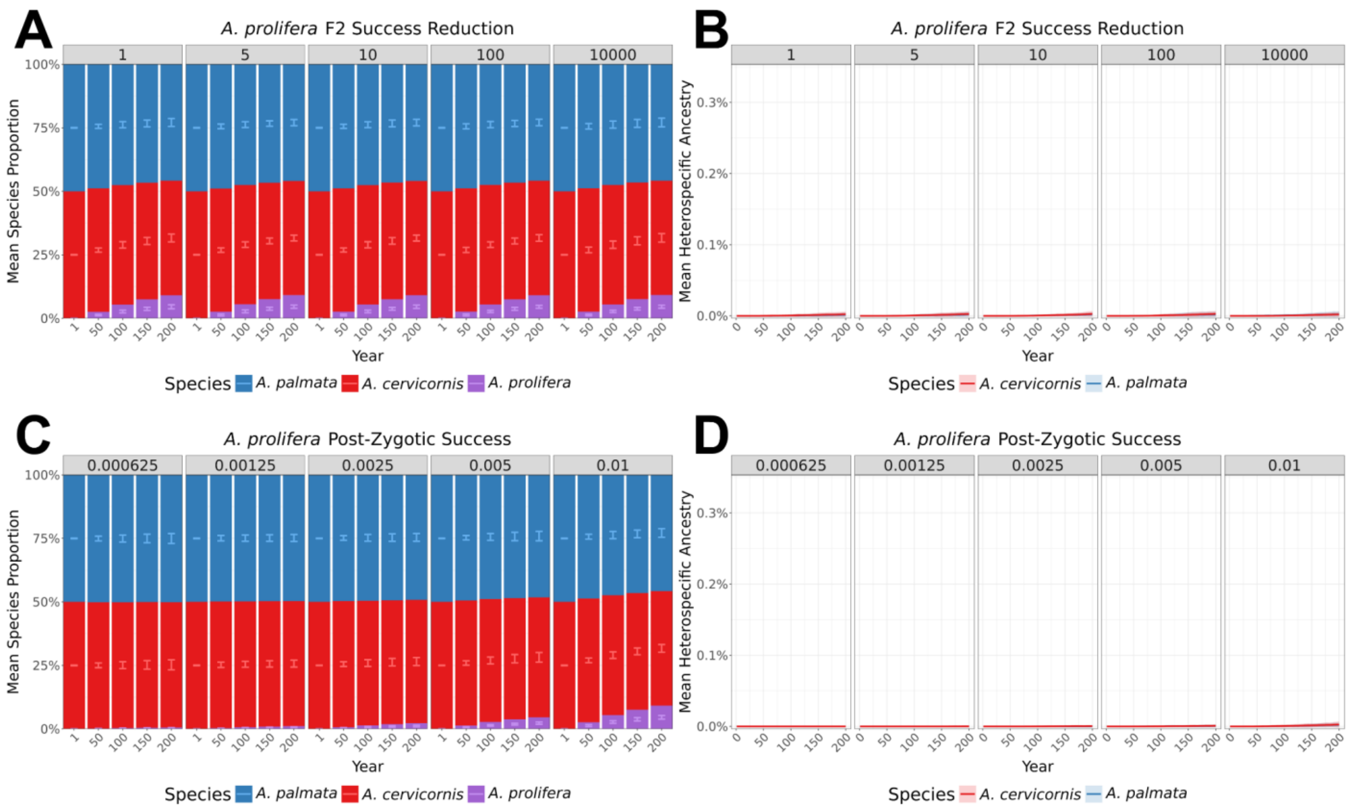


***Supplementary Figure 4:*** **Effect of $F_2$ success reduction and post-zygotic isolation on introgression and species dynamics within first 200 years**

Compared to Main Text Fig. 2E–H, this plot is the same data but instead focusing on the first 200 years of the simulation (i.e., a shorter timespan than Main Text Fig. 2E–H). Species ratios (A, C) and mean heterospecific ancestry (B, D) through time faceted by $F_2$ success reduction with 5×, 10×, 100×, or 1,000× relative to the baseline initial larval survival (A, B), and *A. prolifera* larval success with halved survival in four steps starting from our baseline 1% to 0.5%, 0.025%, 0.0125%, 0.00625% (C, D). For species-ratio panels (A, C), the x-axis shows simulation years and the y-axis the mean species ratio across replicates (n=50). Stacked bars display the mean proportion of each taxon, with error bars indicating ±2× standard deviation between replicates (n=50). Colors: blue = *A. palmata*, red = *A. cervicornis*, purple = hybrid *A. prolifera*. For heterospecific-ancestry panels (B, D), solid lines represent mean heterospecific ancestry across replicates (n=50) and shaded areas show ±2× the standard deviation between replicates. Colors: blue = *A. palmata*, red = *A. cervicornis*.

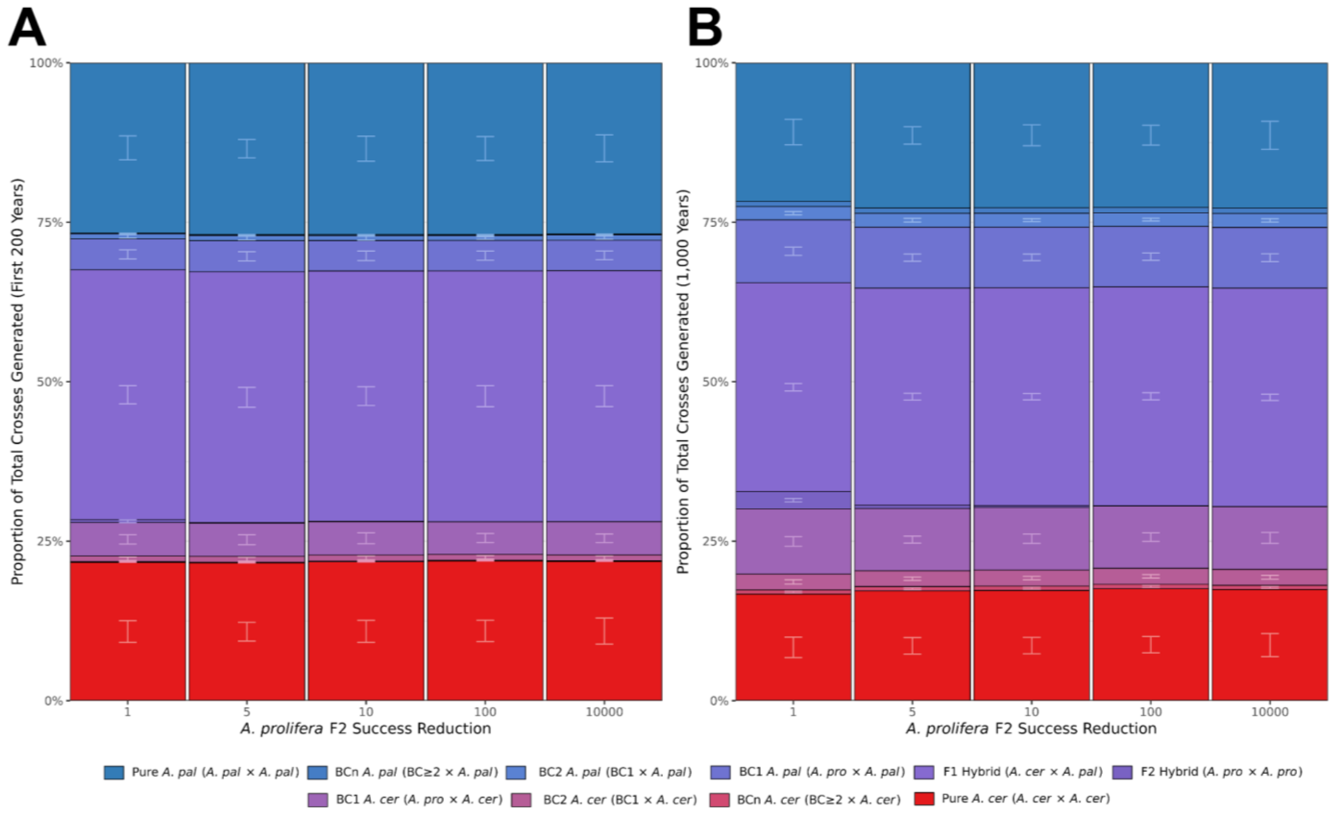


***Supplementary Figure 5:*** **Parentage proportions of crosses under different amounts of $F_2$ success reduction**

Unlike the standard species dynamic ratio plots, the x-axis of this figure represents the $F_2$ success reduction rather than the simulation year. Additionally, compared to standard species dynamic plots, instead of there being three categories (with blue bars representing *A. palmata*, red bars representing *A. cervicornis*, and purple bars representing *A. prolifera*), these stacked bars break out the hybrid *A. prolifera* bar into four different bars representing $F_1$s, $F_2$s, $BC_1$s and $BC_2$s. The parental bars are broken into two separate categories, $BC_3$+ and pure parentals with no history of backcrossing. The stacked bars represent a mean of the total proportion of the respective crosses across 50 replicates. The error bars represent ±2× the standard deviation between replicates. (A) Shows the total parentage proportions generated in the first 200 years of the simulation. (B) Shows the total parentage proportions generated over all 1,000 years of the simulation.

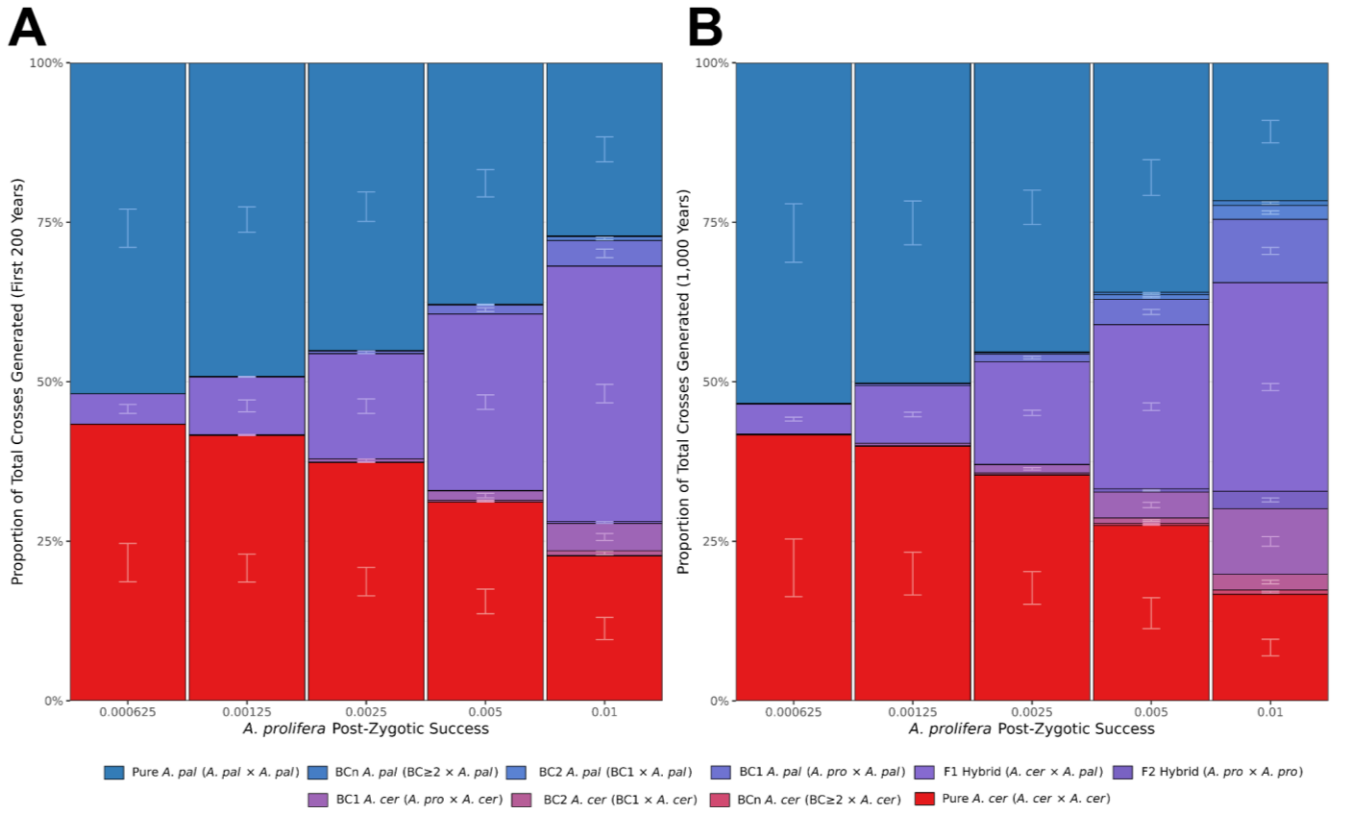


***Supplementary Figure 6:*** **Parentage proportions of crosses under different amounts of *A. prolifera* post-zygotic success**

Unlike the standard species dynamic ratio plots, the x-axis of this figure represents values of *A. prolifera* larval success rates rather than the simulation year. With halved survival in five steps from our baseline of 1% to 0.5%, 0.025%, 0.0125% and 0.00625%. Additionally, compared to standard species dynamic plots, instead of there being three categories (with blue bars representing *A. palmata*, red bars representing *A. cervicornis*, and purple bars representing *A. prolifera*), these stacked bars break out the hybrid *A. prolifera* bar into four different bars representing $F_1$s, $F_2$s, $BC_1$s and $BC_2$s. The parental bars are broken into two separate categories, $BC_3$+ and pure parentals with no history of backcrossing. The stacked bars represent a mean of the total proportion of the respective crosses across 50 replicates. The error bars represent ±2× the standard deviation between replicates. (A) Shows the total parentage proportions generated in the first 200 years of the simulation. (B) Shows the total parentage proportions generated over all 1,000 years of the simulation.

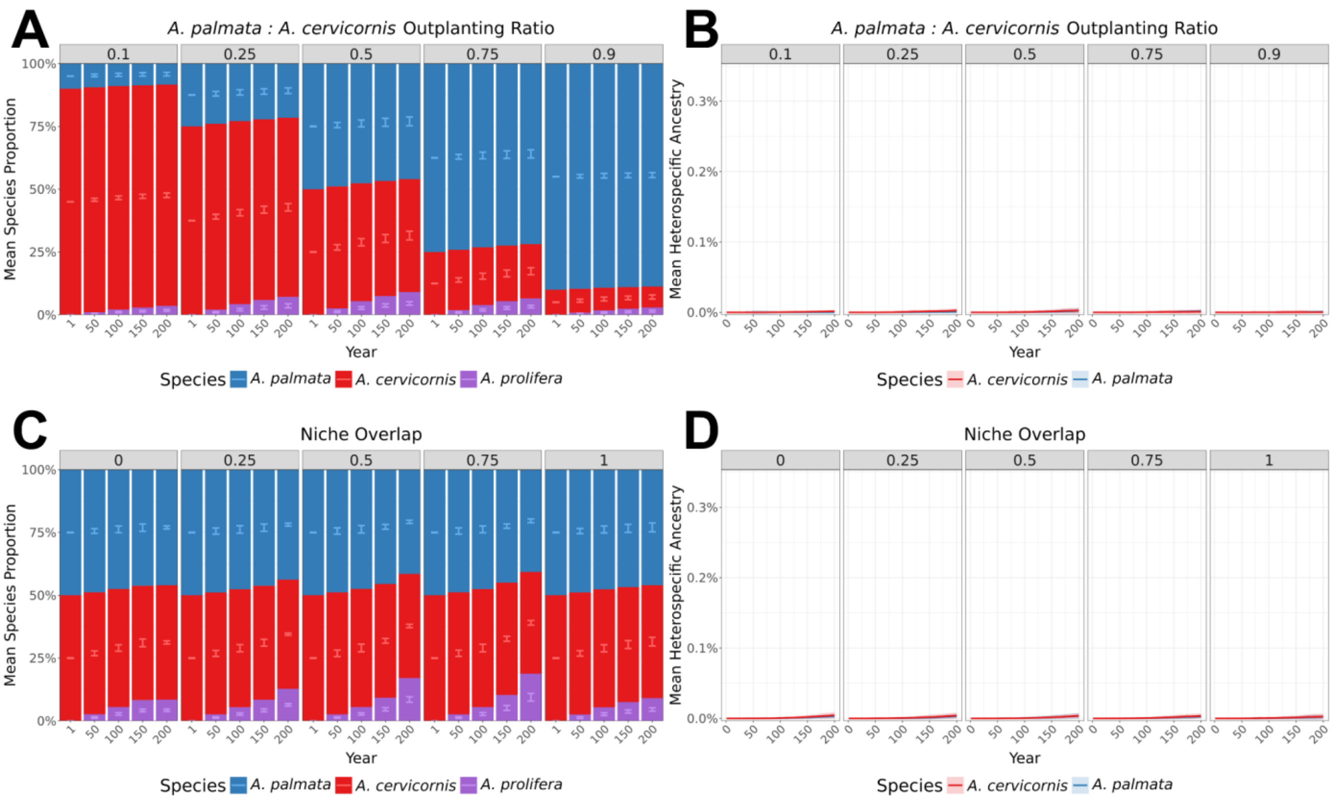


***Supplementary Figure 7:*** **Effect of initial species ratios and niche overlap on introgression and species dynamics within first 200 years**

Compared to Main Text Fig. 3, this plot is the same data but instead focusing on the first 200 years of the simulation (i.e., a shorter timespan than Main Text Fig. 3). Species ratios (A, C) and mean heterospecific ancestry (B, D) through time under two conditions: varying initial outplanting ratios of *A. palmata* to *A. cervicornis* where a ratio of 0.1 means 10%:90% *A. palmata:A. cervicornis* and 0.9 means 90%:10% *A. palmata:A. cervicornis* (A, B) and varying degrees of niche overlap where 0 = no overlap and 1 = full overlap (C, D). For species-ratio panels (A, C), the x-axis shows simulation years and the y-axis the mean species ratio across replicates (n=50). Stacked bars display the mean proportion of each taxon, with error bars indicating ±2× standard deviation between replicates. Colors: blue = *A. palmata*, red = *A. cervicornis*, purple = hybrid *A. prolifera*. For heterospecific-ancestry panels (B, D), solid lines represent mean heterospecific ancestry across replicates (n=50) and shaded areas show ±2× the standard deviation between replicates. Colors: blue = *A. palmata*, red = *A. cervicornis*.

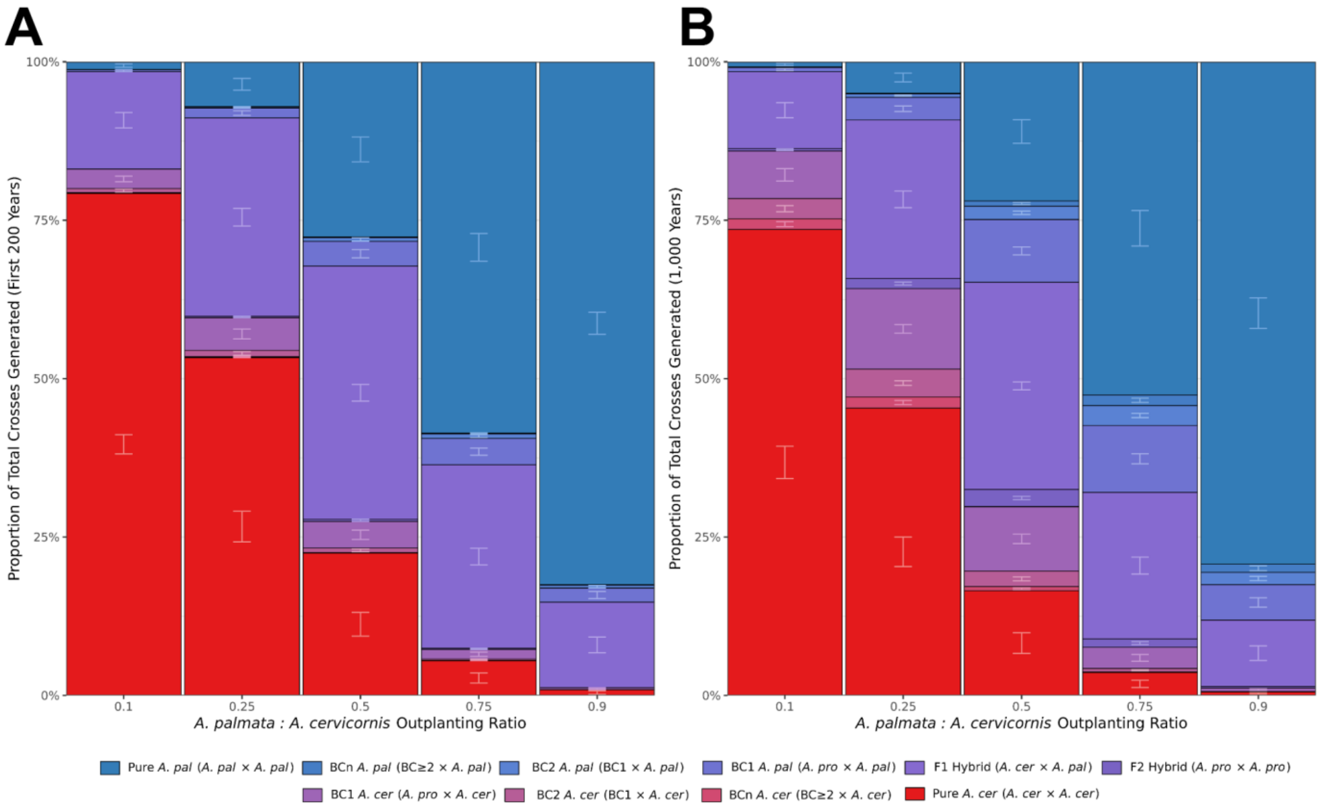


***Supplementary Figure 8:*** **Parentage proportions of crosses under different initial species ratios** Unlike the standard species dynamic ratio plots, the x-axis of this figure represents the initial species ratios where a ratio of 0.1 means 10%:90% *A. palmata:A. cervicornis* and 0.9 means 90%:10% *A. palmata:A. cervicornis,* rather than the simulation year. Additionally, compared to standard species dynamic plots, instead of there being three categories (with blue bars representing *A. palmata*, red bars representing *A. cervicornis*, and purple bars representing *A. prolifera*), these stacked bars break out the hybrid *A. prolifera* bar into four different bars representing $F_1$s, $F_2$s, $BC_1$s and $BC_2$s. The parental bars are broken into two separate categories, $BC_3$+ and pure parentals with no history of backcrossing. The stacked bars represent a mean of the total proportion of the respective crosses across 50 replicates. The error bars represent ±2× the standard deviation between replicates. (A) Shows the total parentage proportions generated in the first 200 years of the simulation. (B) Shows the total parentage proportions generated over all 1,000 years of the simulation.

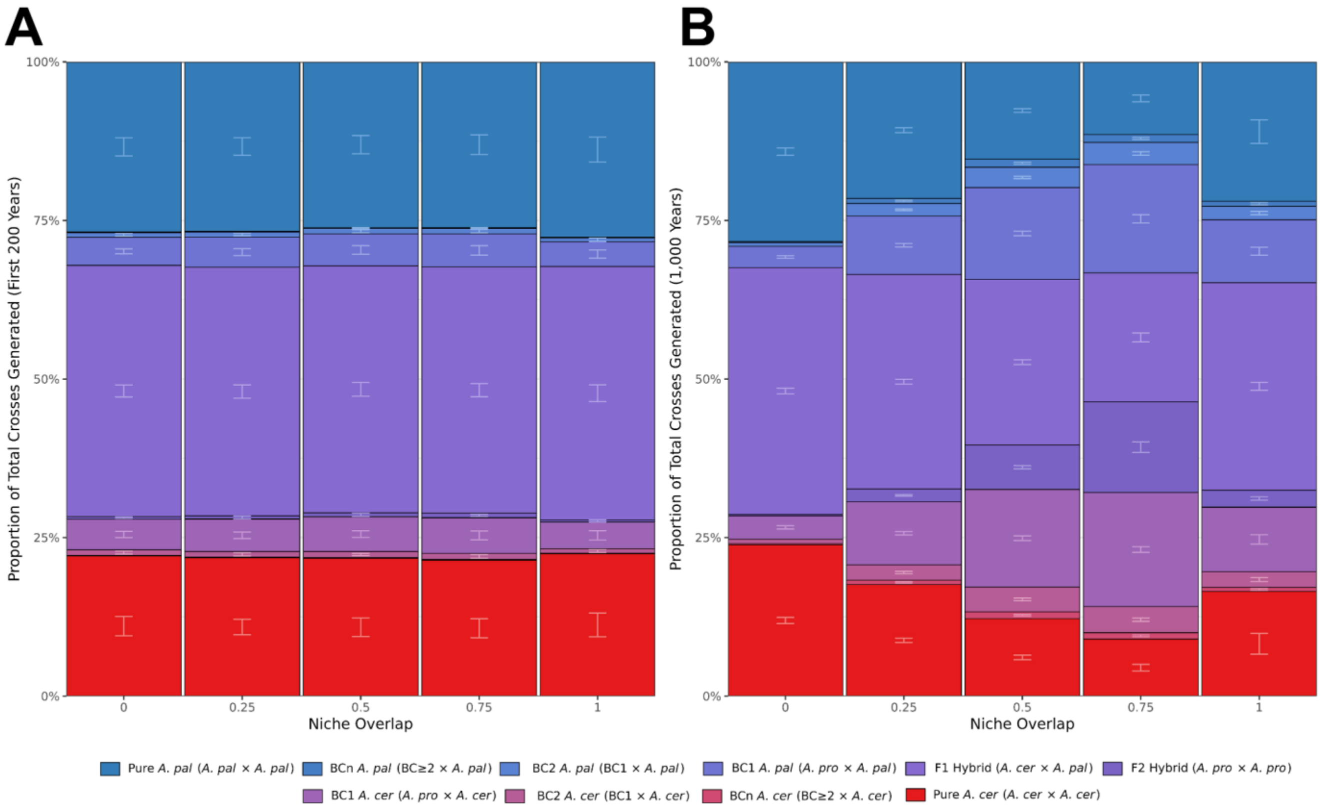


***Supplementary Figure 9:*** **Parentage proportions of crosses under different niche overlap values** Unlike the standard species dynamic ratio plots, the x-axis of this figure represents the niche overlap values where 0 = no overlap and 1 = full overlap, rather than the simulation year. Additionally, compared to standard species dynamic plots, instead of there being three categories (with blue bars representing *A. palmata*, red bars representing *A. cervicornis*, and purple bars representing *A. prolifera*), these stacked bars break out the hybrid *A. prolifera* bar into four different bars representing $F_1$s, $F_2$s, $BC_1$s and $BC_2$s. The parental bars are broken into two separate categories, $BC_3$+ and pure parentals with no history of backcrossing. The stacked bars represent a mean of the total proportion of the respective crosses across 50 replicates. The error bars represent ±2× the standard deviation between replicates. (A) Shows the total parentage proportions generated in the first 200 years of the simulation. (B) Shows the total parentage proportions generated over all 1,000 years of the simulation.

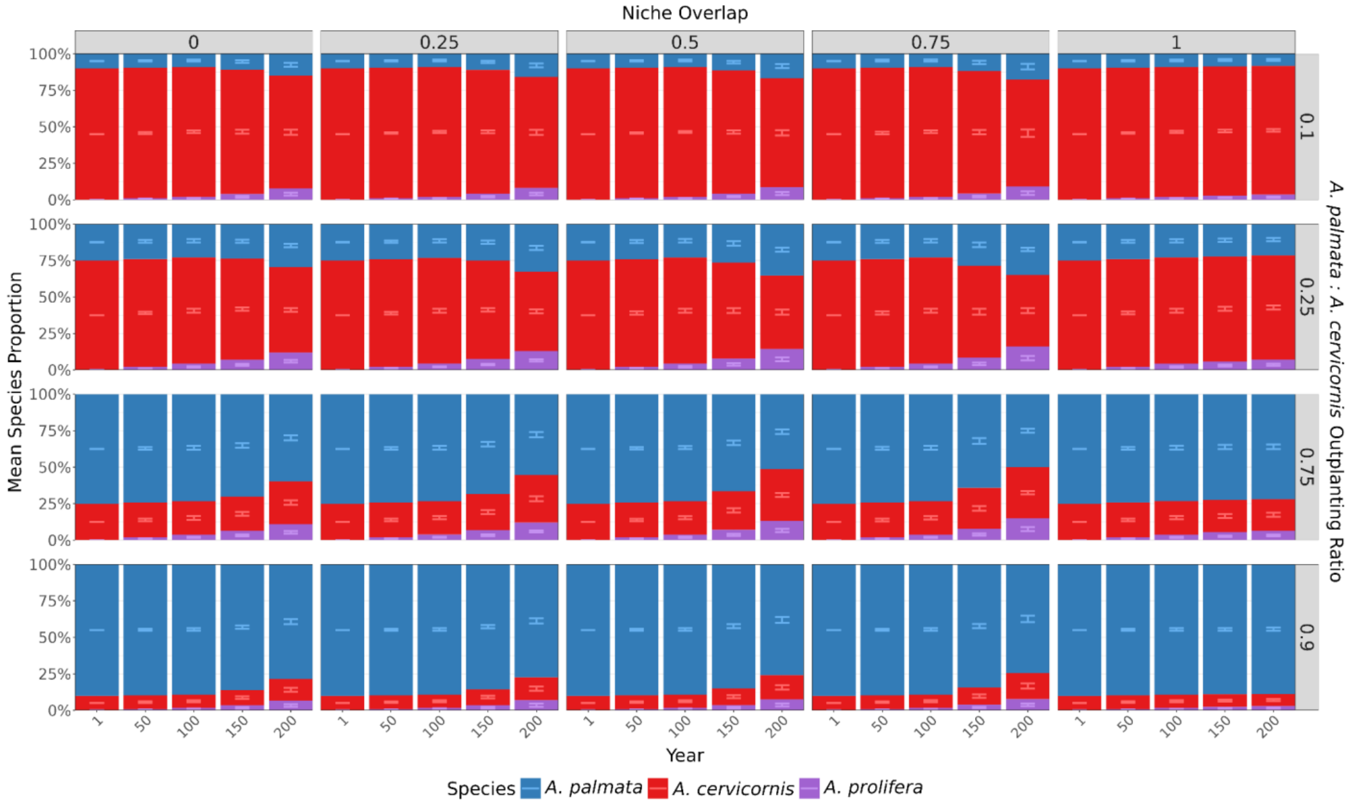


***Supplementary Figure 10:*** **Effect of the interaction between initial outplanting ratio and niche overlap on species dynamics within first 200 years**

Compared to Main Text Fig. 4, this plot is the same data but instead focusing on the first 200 years of the simulation (i.e., a shorter timespan than Main Text Fig. 4). Species ratios through time, faceted the degree of niche overlap where 0 = no overlap and 1 = full overlap (top), and the initial outplanting ratio of *A. palmata* to *A. cervicornis* (right) where a ratio of 0.1 means 10%:90% *A. palmata:A. cervicornis* and 0.9 means 90%:10% *A. palmata:A. cervicornis*. The x-axis represents snapshots of the mean species ratio (y-axis) in different years through the simulation. The stacked bars represent a mean of the species ratio in the population across replicates. The error bars represent ±2× the standard deviation between replicates (n=50). Blue bars represent *A. palmata*, red bars represent *A. cervicornis*, and purple bars represent their hybrid, A. prolifera.

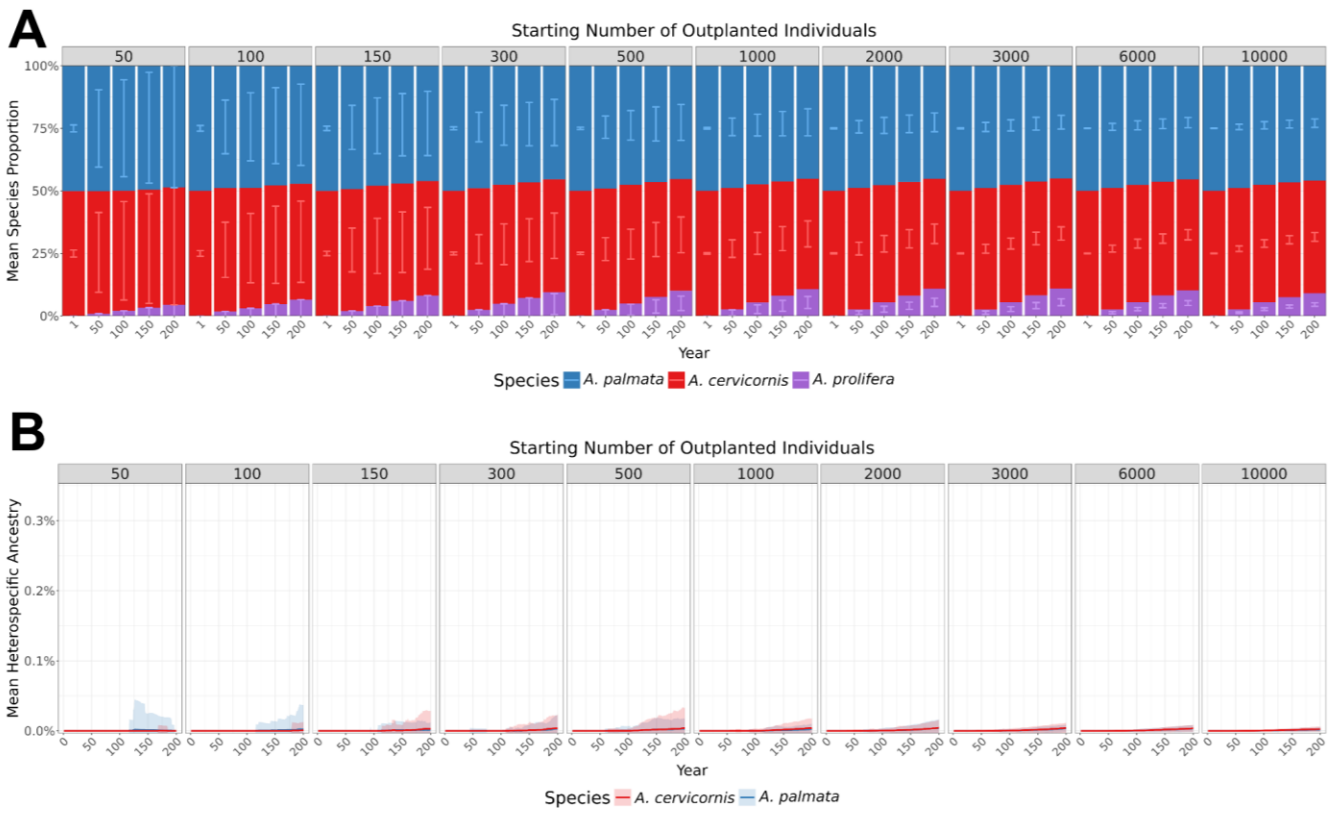


***Supplementary Figure 11:*** **Effect of outplant project size on restoration outcomes within first 200 years**

Compared to Main Text Fig. 5, this plot is the same data but instead focusing on the first 200 years of the simulation (i.e., a shorter timespan than Main Text Fig. 5). (A) Species ratios through time, faceted the number of initially outplanted individuals ranging from 50–10,000 outplanted coral fragments. X-axis represents snapshots of the mean species ratio (y-axis) in different years through the simulation. The stacked bars represent a mean of the species ratio in the population across replicates. The error bars represent ±2× the standard deviation between replicates (n=150). Blue bars represent *A. palmata*, red bars represent *A. cervicornis*, and purple bars represent their hybrid, *A. prolifera*. (B) Mean heterospecific ancestry (i.e., the amount of ancestry from the other parental species) through time, faceted by the number of initially outplanted individuals. The solid lines represent the mean heterospecific ancestry in the population across replicates and shaded areas show ±2× the standard deviation between replicates (n=150). Blue lines and shading represent *A. palmata*, red lines and shading represent *A. cervicornis*.

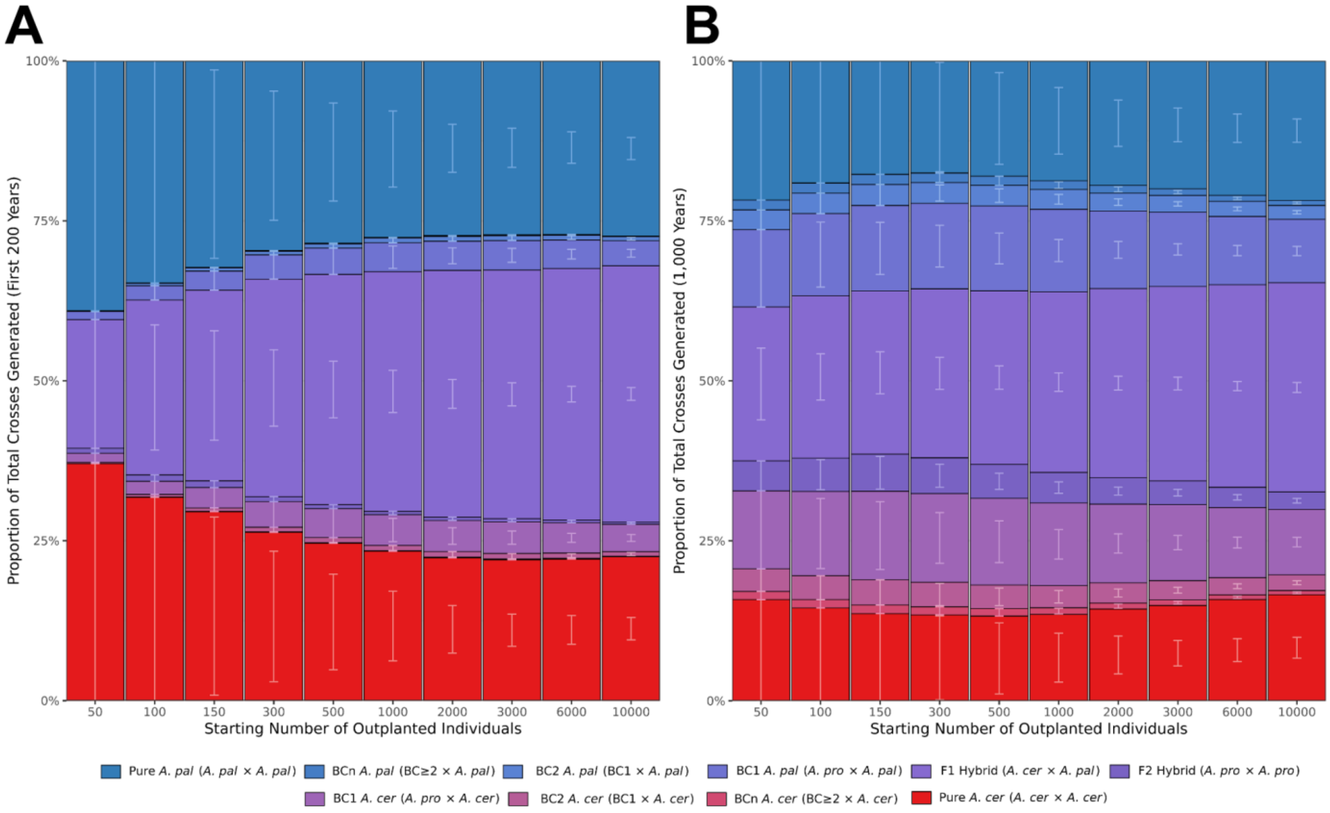


***Supplementary Figure 12:*** **Parentage proportions of crosses under different numbers of initially outplanted individuals**

Unlike the standard species dynamic ratio plots, the x-axis of this figure represents the starting number of outplanted individuals rather than the simulation year. Additionally, compared to standard species dynamic plots, instead of there being three categories (with blue bars representing *A. palmata*, red bars representing *A. cervicornis*, and purple bars representing *A. prolifera*), these stacked bars break out the hybrid *A. prolifera* bar into four different bars representing $F_1$s, $F_2$s, $BC_1$s and $BC_2$s. The parental bars are broken into two separate categories, $BC_3$+ and pure parentals with no history of backcrossing. The stacked bars represent a mean of the total proportion of the respective crosses across 50 replicates. The error bars represent ±2× the standard deviation between replicates. (A) Shows the total parentage proportions generated in the first 200 years of the simulation. (B) Shows the total parentage proportions generated over all 1,000 years of the simulation.

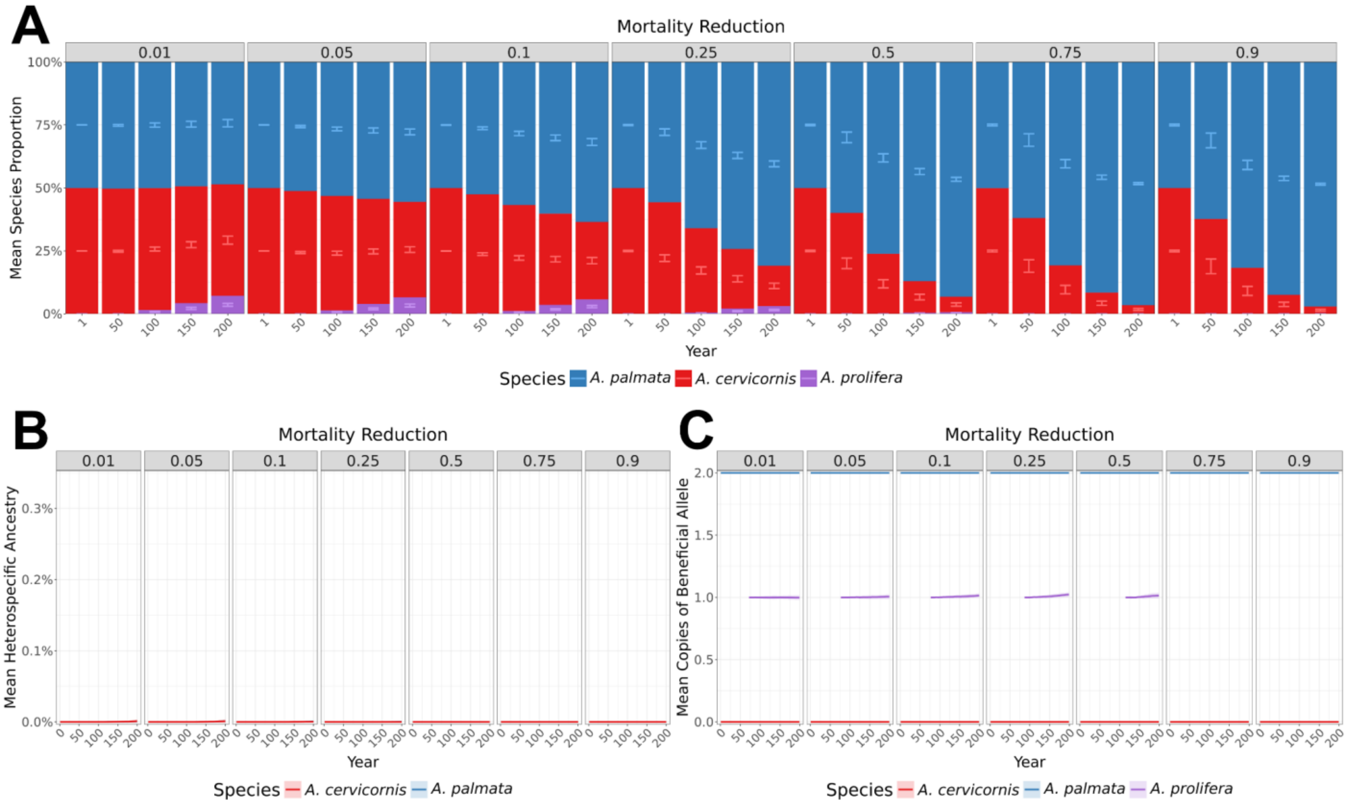


***Supplementary Figure 13:*** **Effect of single beneficial loci initially fixed in A. palmata on introgression and species dynamics within first 200 years**

Compared to Main Text Fig. 6, this plot is the same data but instead focusing on the first 200 years of the simulation (i.e., a shorter timespan than Main Text Fig. 6). (A) Species ratios through time, faceted the strength of the beneficial allele in *A. palmata*. X-axis represents snapshots of the mean species ratio (y-axis) in different years through the simulation ranging from 1%–90% mortality reduction. The stacked bars represent a mean of the species ratio in the population across replicates. The error bars represent ±2× the standard deviation between replicates (n=50). Blue bars represent *A. palmata*, red bars represent *A. cervicornis*, and purple bars represent their hybrid, A. prolifera*.* (B) Mean heterospecific ancestry (i.e., the amount of ancestry from the other parental species) through time, faceted the mortality reduction from carrying the beneficial allele. The solid lines represent the mean heterospecific ancestry in the population across replicates and shaded areas show ±2× the standard deviation between replicates (n=50). Blue lines and shading represent *A. palmata*, red lines and shading represent *A. cervicornis.* (C) The allele frequency of the beneficial allele in the three Acroporids through time.

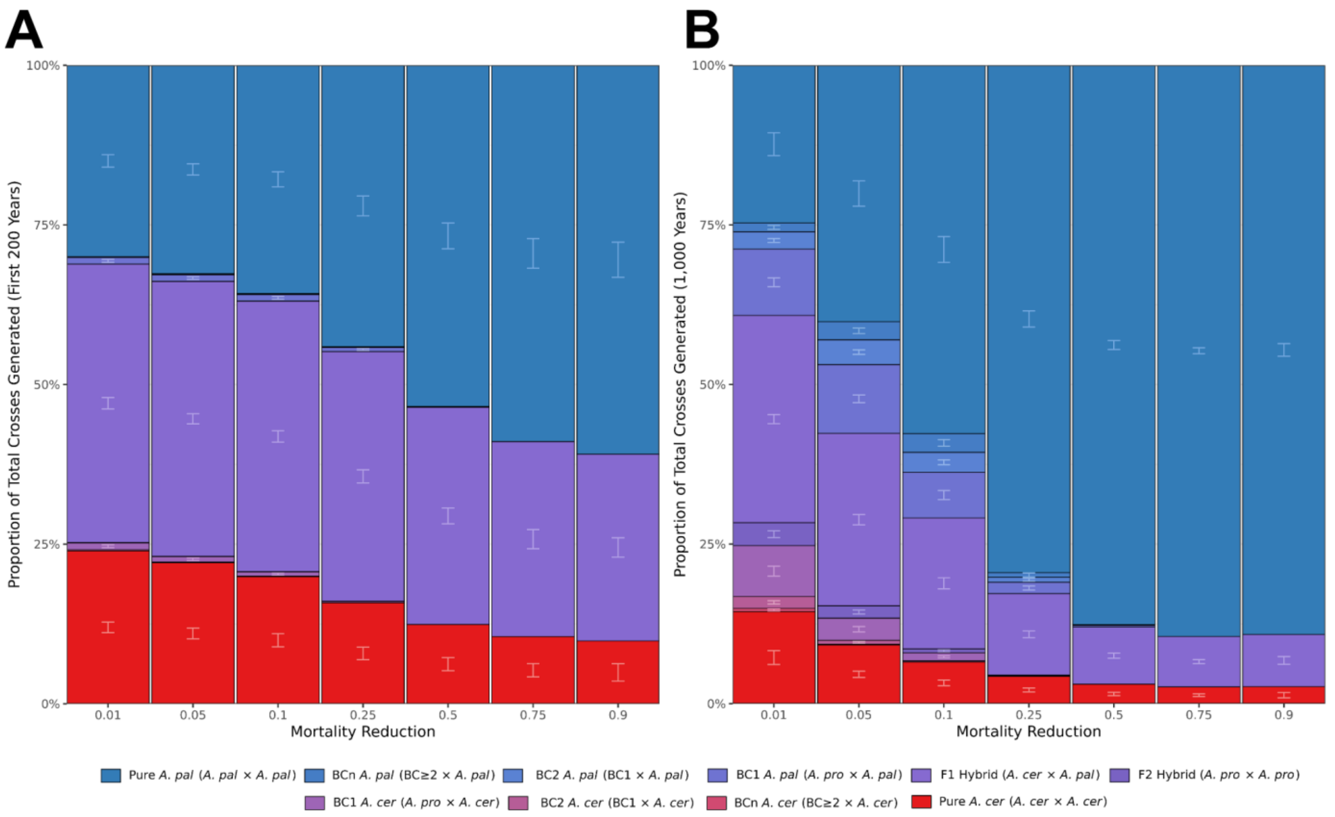


***Supplementary Figure 14:*** **Parentage proportions of crosses under different values of mortality reduction under an additive mutation model**

Unlike the standard species dynamic ratio plots, the x-axis of this figure represents the mortality reduction values rather than the simulation year. Additionally, compared to standard species dynamic plots, instead of there being three categories (with blue bars representing *A. palmata*, red bars representing *A. cervicornis*, and purple bars representing *A. prolifera*), these stacked bars break out the hybrid *A. prolifera* bar into four different bars representing $F_1$s, $F_2$s, $BC_1$s and $BC_2$s. The parental bars are broken into two separate categories, $BC_3$+ and pure parentals with no history of backcrossing. The stacked bars represent a mean of the total proportion of the respective crosses across 50 replicates. The error bars represent ±2× the standard deviation between replicates. (A) Shows the total parentage proportions generated in the first 200 years of the simulation. (B) Shows the total parentage proportions generated over all 1,000 years of the simulation.

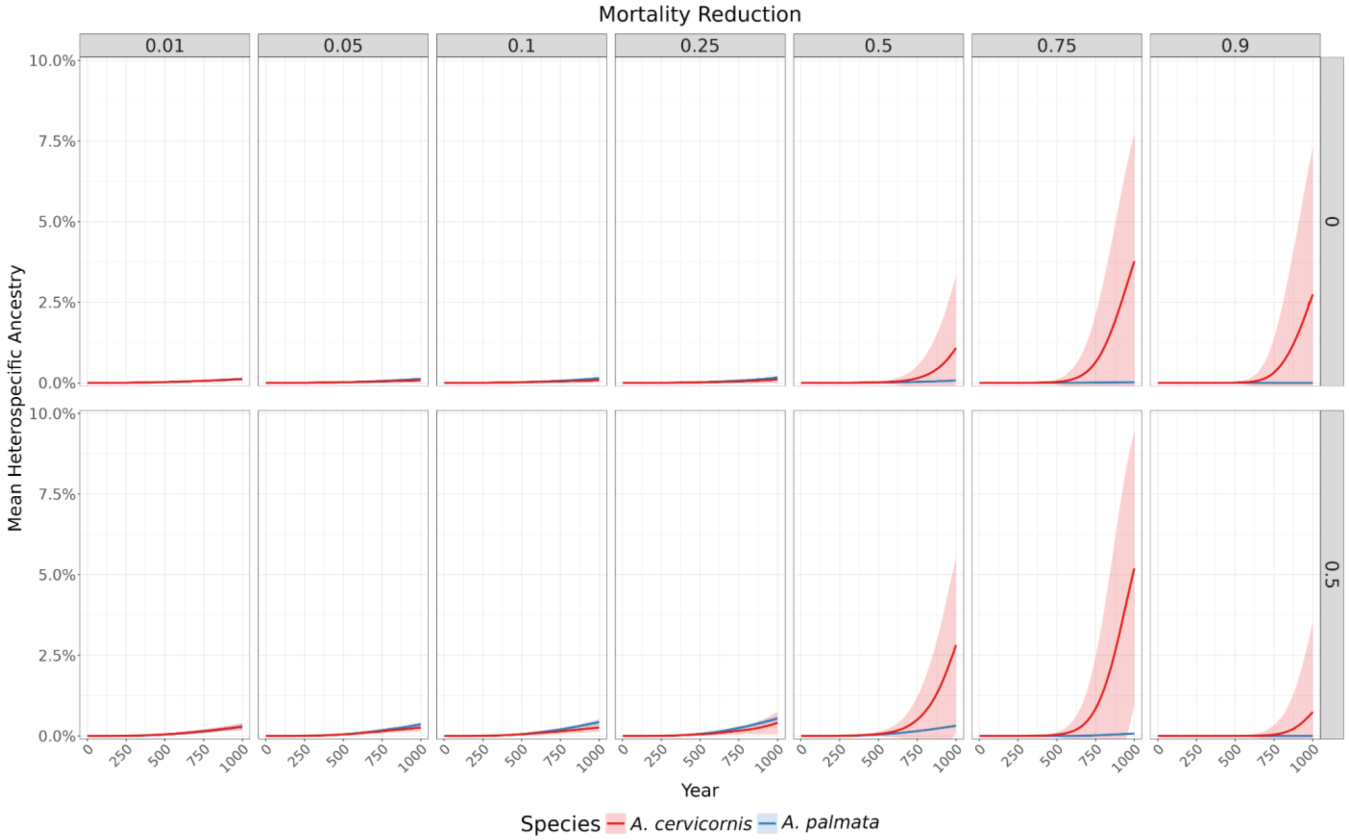


***Supplementary Figure 15:*** **Effect of niche overlaps and single additive beneficial locus initially fixed in A. palmata on heterospecific ancestry transfer**

Mean heterospecific ancestry (i.e., the amount of ancestry from the other parental species) through time, faceted the mortality reduction from carrying the beneficial allele (top) and the niche overlap (right). The solid lines represent the mean heterospecific ancestry in the population across replicates and shaded areas show ±2× the standard deviation between replicates (n=50). Blue lines and shading represent *A. palmata*, red lines and shading represent *A. cervicornis*.

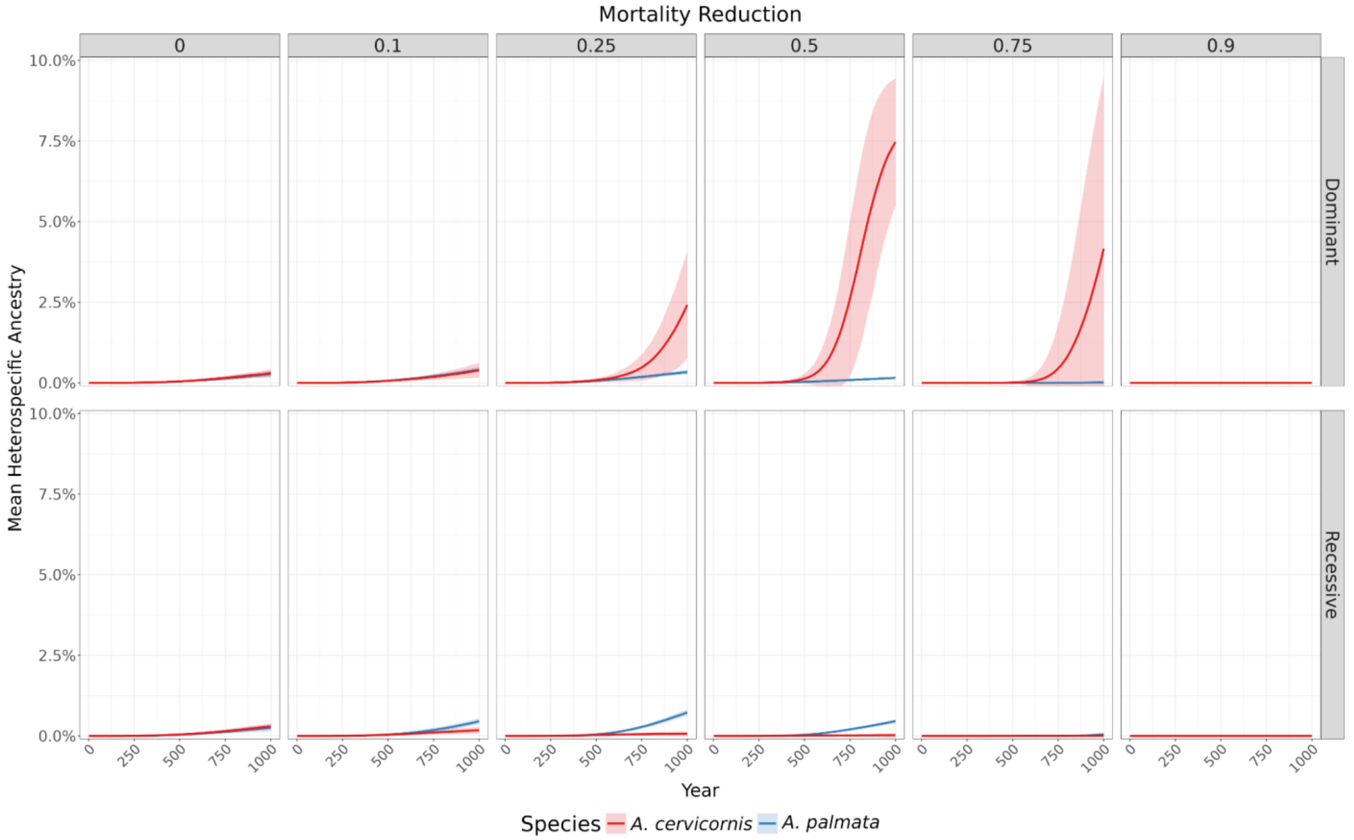


***Supplementary Figure 16:*** **Effect of mutation dominance and single additive beneficial locus initially fixed in *A. palmata* on heterospecific ancestry transfer**

Mean heterospecific ancestry (i.e., the amount of ancestry from the other parental species) through time, faceted the mortality reduction from carrying the beneficial allele (top) and the mutation dominance (right). The solid lines represent the mean heterospecific ancestry in the population across replicates and shaded areas show ±2× the standard deviation between replicates (n=50). Blue lines and shading represent A. palmata, red lines and shading represent A. cervicornis.

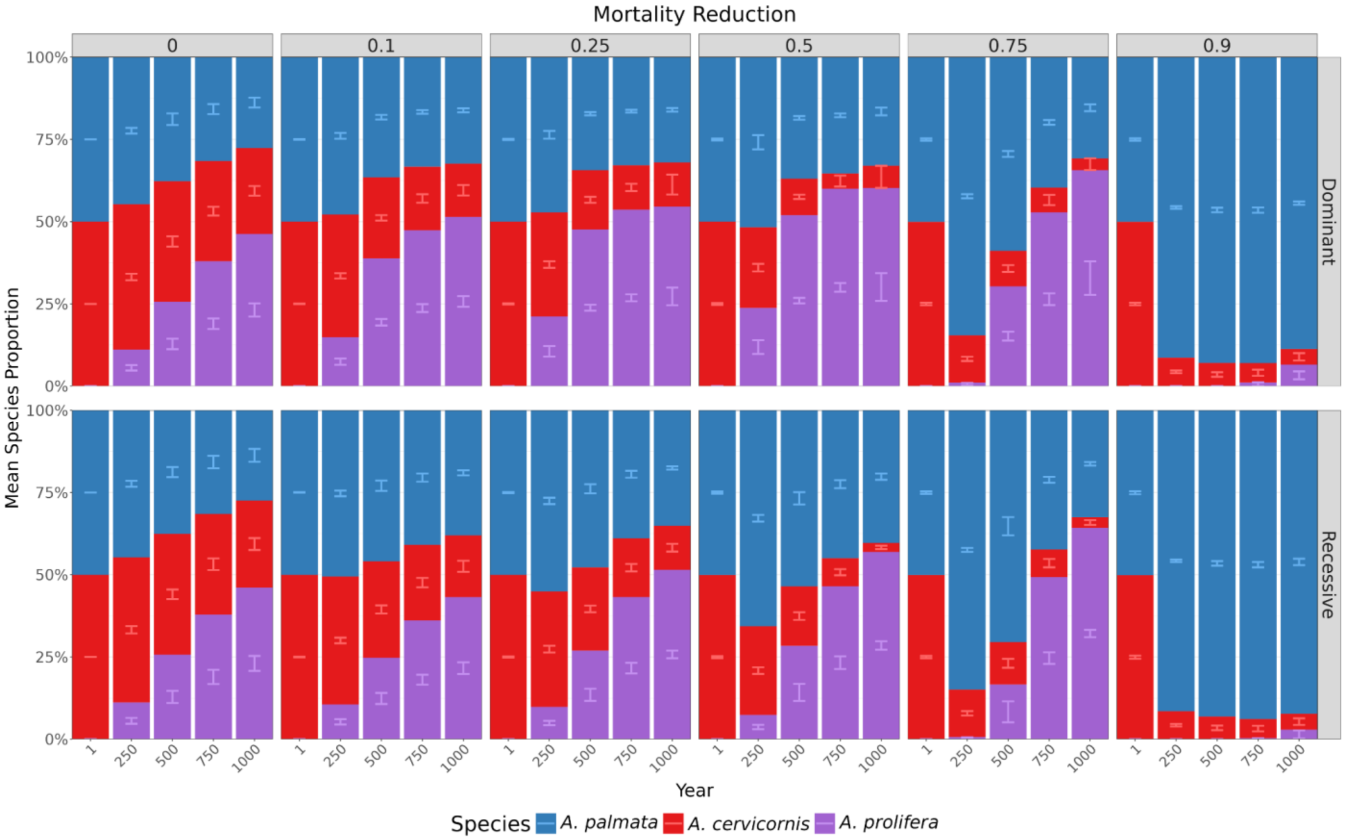


***Supplementary Figure 17:*** **Effect of mutation dominance and single additive beneficial locus initially fixed in *A. palmata* on species dynamics**

Species ratios through time, faceted by the mortality reduction from carrying the beneficial allele (top) and the mutation dominance (right). X-axis represents snapshots of the mean species ratio (y-axis) in different years through the simulation. The stacked bars represent a mean of the species ratio in the population across replicates. The error bars represent ±2× the standard deviation between replicates (n=50). Blue bars represent *A. palmata*, red bars represent *A. cervicornis*, and purple bars represent their hybrid, *A. prolifera.*

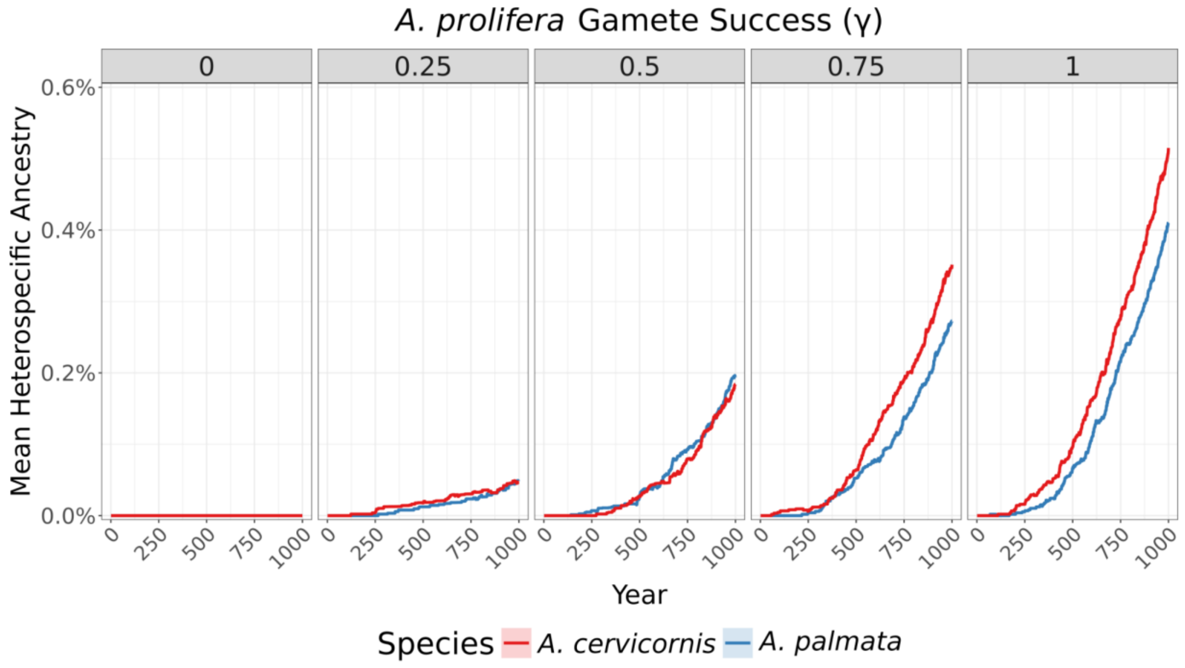


***Supplementary Figure 18:*** **Effect of hybrid gamete success (γ) on introgression without clonal reproduction**

Mean heterospecific ancestry (i.e., the amount of ancestry from the other parental species) through time, faceted by hybrid gamete success (γ), under sims that do not simulate clonal reproduction. The solid lines represent the mean heterospecific ancestry in the population. Blue lines and shading represent *A. palmata*, red lines and shading represent *A. cervicornis*.

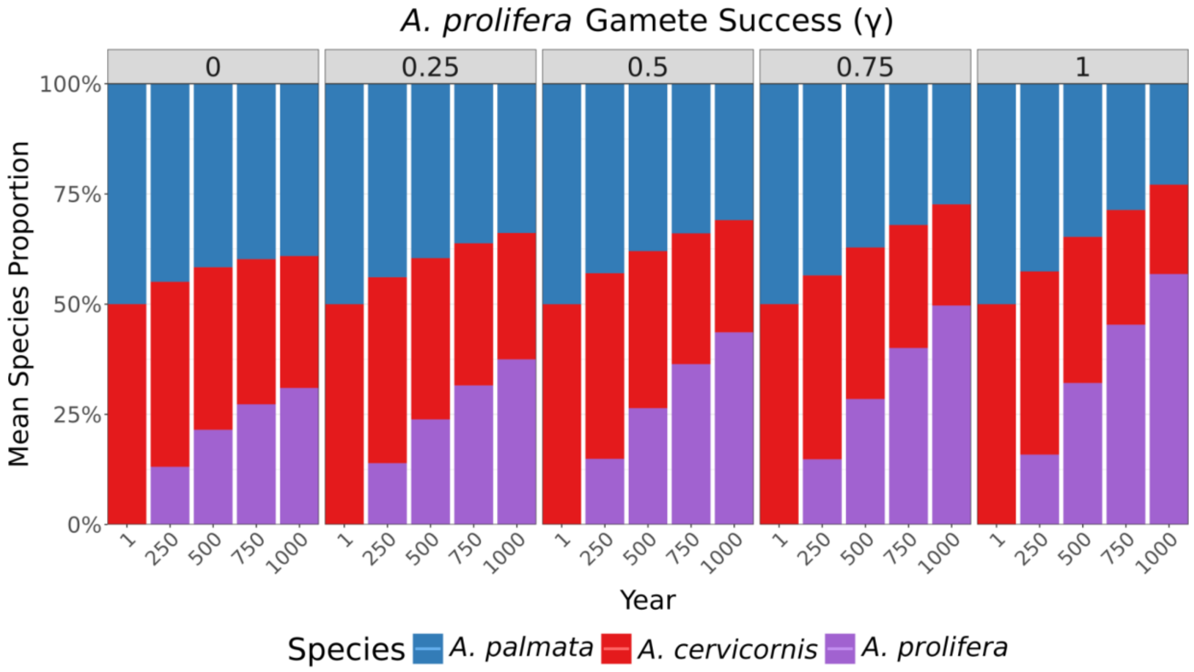


***Supplementary Figure 19:* Effect of hybrid gamete success (γ) on species dynamics without clonal reproduction**

Species ratios through time, faceted by hybrid gamete success (γ), under sims that do not simulate clonal reproduction. X-axis represents snapshots of the mean species ratio (y-axis) in different years through the simulation. The stacked bars represent a mean of the species ratio in the population across replicates. Blue bars represent *A. palmata*, red bars represent *A. cervicornis*, and purple bars represent their hybrid, *A. prolifera.*

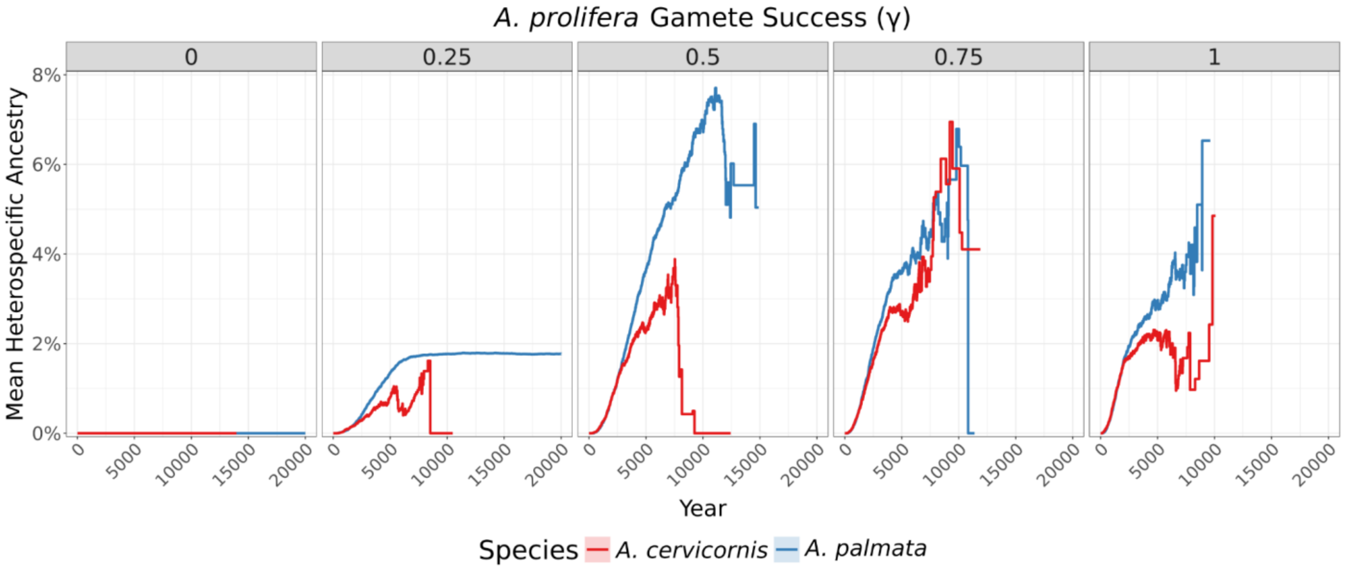


***Supplementary Figure 20:*** **Effect of hybrid gamete success (γ) on long term introgression (20,000 years) without clonal reproduction**

Mean heterospecific ancestry through time, faceted by hybrid gamete success (γ), under sims that do not simulate clonal reproduction and run for 20,000 years. The solid lines represent the mean heterospecific ancestry in the population. Blue lines and shading represent *A. palmata*, red lines and shading represent *A. cervicornis*.

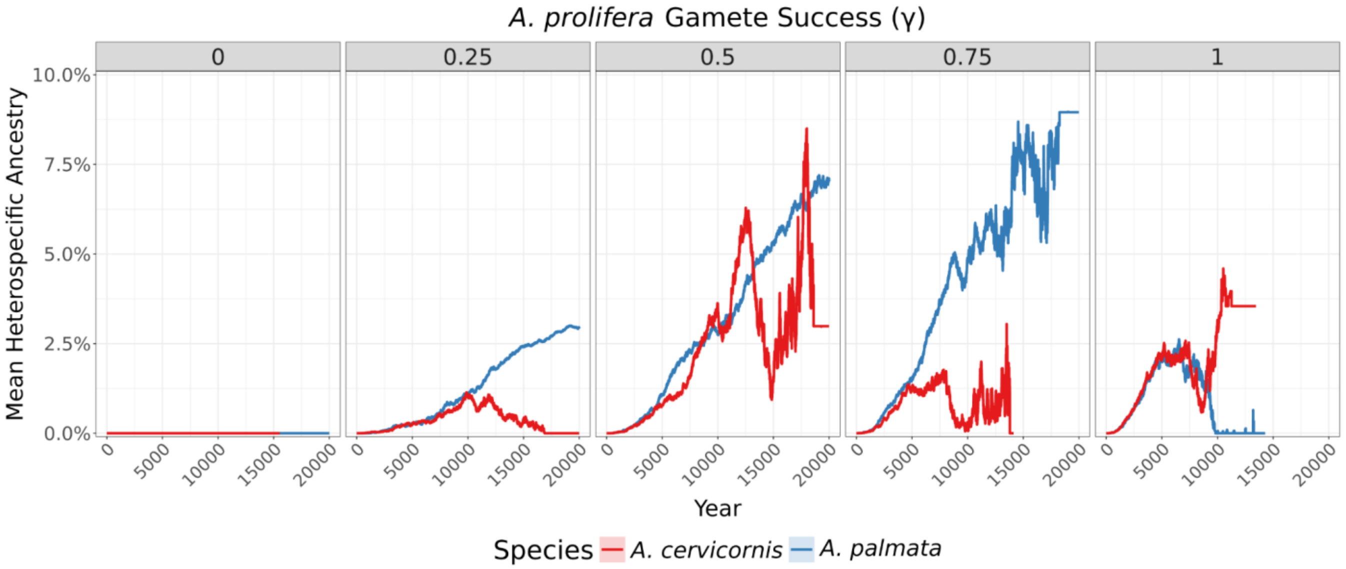


***Supplementary Figure 21:*** **Effect of hybrid gamete success (γ) on long term introgression (20,000 years)**

Mean heterospecific ancestry through time, faceted by hybrid gamete success (γ), run for 20,000 years. The solid lines represent the mean heterospecific ancestry in the population. Blue lines and shading represent *A. palmata*, red lines and shading represent *A. cervicornis*.

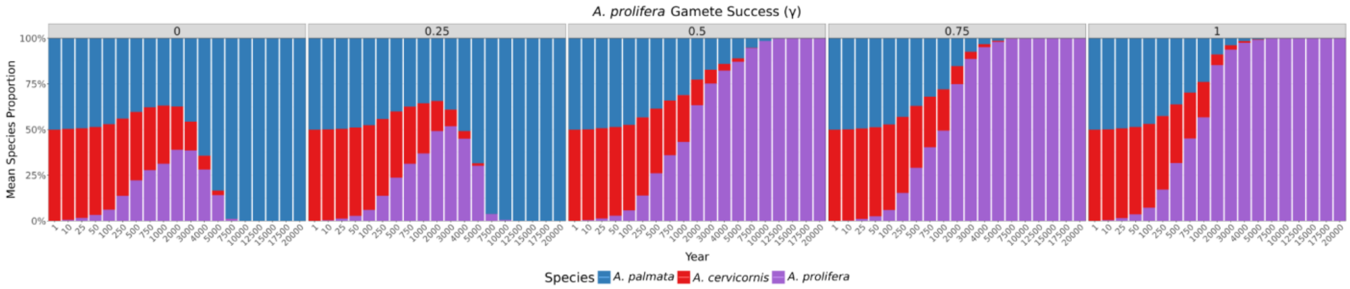


***Supplementary Figure 22:*** **Effect of hybrid gamete success (γ) on long term species dynamics (20,000 years) without clonal reproduction**

Species ratios through time, faceted by hybrid gamete success (γ), under sims that do not simulate clonal reproduction and run for 20,000 years. X-axis represents snapshots of the mean species ratio (y-axis) in different years through the simulation. The stacked bars represent a mean of the species ratio in the population across replicates. Blue bars represent A. palmata, red bars represent A. cervicornis, and purple bars represent their hybrid, A. prolifera.

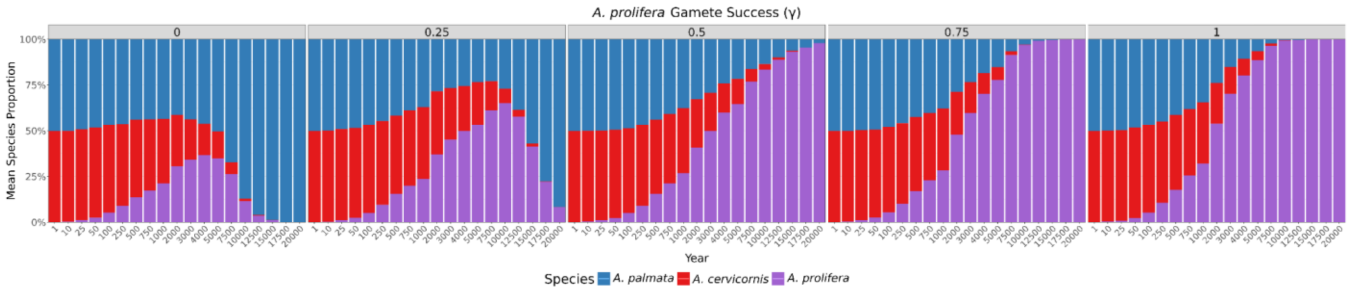


***Supplementary Figure 23: Effect of hybrid gamete success (γ) on long term species dynamics (20,000 years)***

Species ratios through time, faceted by hybrid gamete success (γ), under sims ran longer than our typical 1,000 year simulation (Main Text Fig. 2A), running instead for 20,000 years. X-axis represents snapshots of the mean species ratio (y-axis) in different years through the simulation. The stacked bars represent a mean of the species ratio in the population across replicates. Blue bars represent *A. palmata*, red bars represent *A. cervicornis*, and purple bars represent their hybrid, *A. prolifera.*

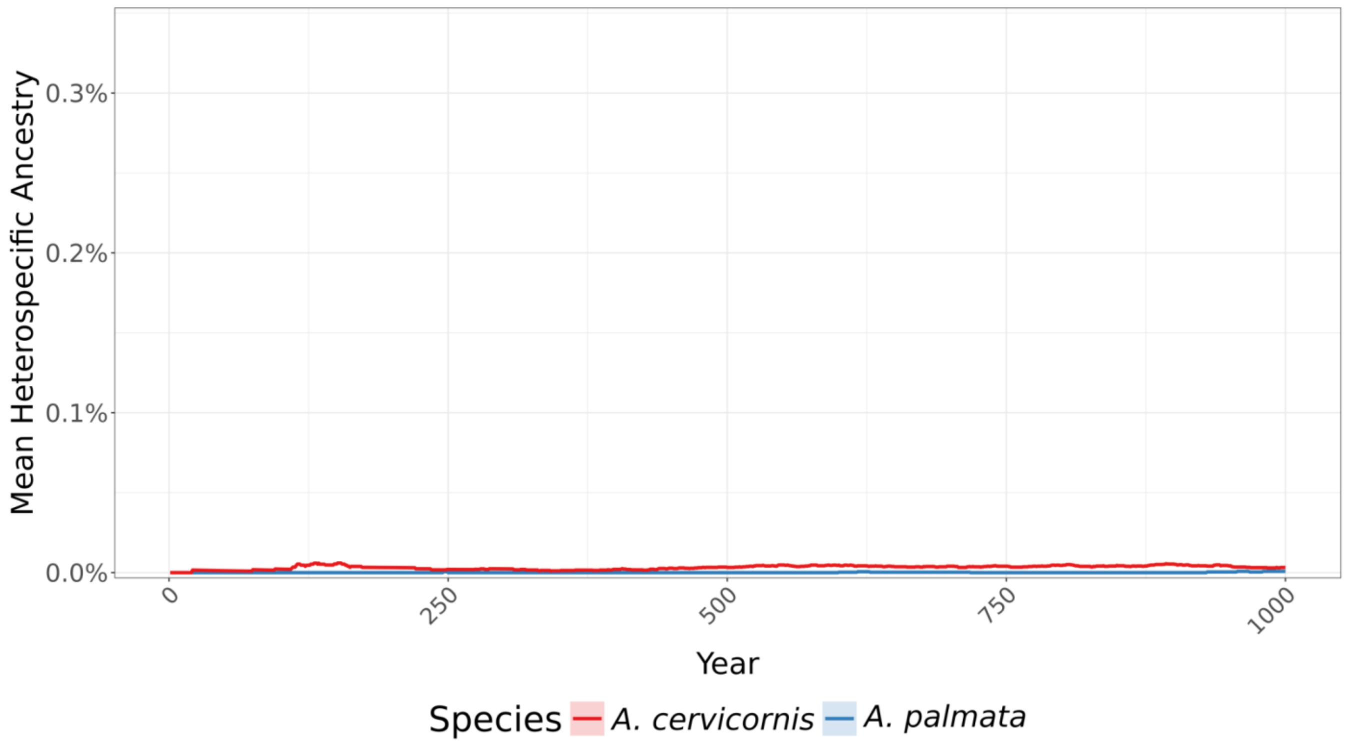


***Supplementary Figure 24:*** **Effect of smaller clonal dispersal distance on introgression levels**
Mean heterospecific ancestry through time, under our baseline parameter combinations, under sims that have reduced the fragment dispersal scale by an order of magnitude (i.e., 10× smaller than the baseline). The solid lines represent the mean heterospecific ancestry in the population. Blue lines and shading represent *A. palmata*, red lines and shading represent *A. cervicornis*.

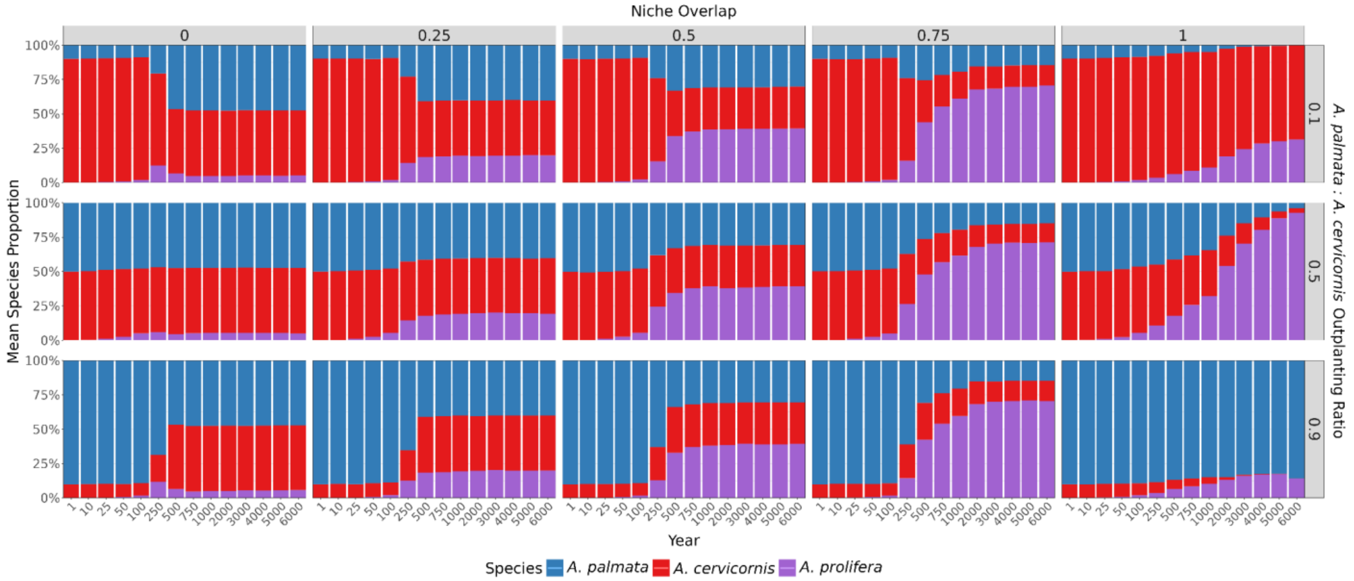


***Supplementary Figure 25:*** **Effect of the interaction between initial outplanting ratio and niche overlap on long term species dynamics (6,000 years)**

Species ratios through time, faceted the degree of niche overlap (top) and the initial outplanting ratio of *A. palmata* to *A. cervicornis* (right), under sims ran longer than our typical 1,000 year simulation (Main Text Fig. 4), running instead for 6,000 years. X-axis represents snapshots of the mean species ratio (y-axis) in different years through the simulation. The stacked bars represent a mean of the species ratio in the population across replicates. Blue bars represent *A. palmata*, red bars represent *A. cervicornis*, and purple bars represent their hybrid, *A. prolifera.*
